# Pharmacological characterization and radiolabeling of VUF15485, a high-affinity small-molecule agonist for the atypical chemokine receptor ACKR3

**DOI:** 10.1101/2023.07.12.548622

**Authors:** Aurelien M. Zarca, Ilze Adlere, Cristina P. Viciano, Marta Arimont-Segura, Max Meyrath, Icaro A. Simon, Jan Paul Bebelman, Dennis Laan, Hans G. J. Custers, Elwin Janssen, Kobus L. Versteegh, Reggie Bosma, Iwan J. P. de Esch, Henry F. Vischer, Maikel Wijtmans, Martyna Szpakowska, Andy Chevigné, Carsten Hoffmann, Chris de Graaf, Barbara A. Zarzycka, Albert D. Windhorst, Martine J. Smit, Rob Leurs

## Abstract

Atypical chemokine receptor 3 (ACKR3), formerly referred to as CXCR7, is considered to be an interesting drug target. In this study we report on the synthesis, pharmacological characterization and radiolabeling of VUF15485, a new ACKR3 small-molecule agonist, that will serve as an important new tool to study this β-arrestin-biased chemokine receptor. VUF15485 binds with nanomolar affinity (pIC_50_ = 8.3) to human ACKR3, as measured in [^125^I]CXCL12 competition binding experiments. Moreover, in a BRET-based β-arrestin2 recruitment assay VUF15485 acts as an ACKR3 agonist with high potency (pEC_50_ = 7.6) and shows a similar extent of receptor activation compared to CXCL12 when using a newly developed, FRET-based ACKR3 conformational sensor. Moreover, the ACKR3 agonist VUF15485 was tested against a (atypical) chemokine receptor panel (agonist and antagonist mode) and proves to be selective for ACKR3. VUF15485 was subsequently labeled with tritium at one of its methoxy groups affording [^3^H]VUF15485. The small-molecule agonist radioligand binds saturably and with high affinity to human ACKR3 (*K*_d_ = 8.2 nM). [^3^H]VUF15485 shows rapid binding kinetics and consequently a short residence time (RT < 2 min) for its binding to ACKR3. Displacement of [^3^H]VUF15485 binding to membranes of HEK293T cells, transiently expressing ACKR3, with a number of CXCR3, CXCR4 or ACKR3 small-molecule ligands confirmed the ACKR3 profile of the [^3^H]VUF15485 binding site. Interestingly, the chemokine ligands CXCL11 and CXCL12 are not able to displace the binding of [^3^H]VUF15485 to ACKR3. The radiolabeled VUF15485 was subsequently used to evaluate its binding pocket. Site-directed mutagenesis and docking studies using a recently solved cryo-EM structure propose VUF15485 to bind in the major and the minor binding pocket of ACKR3.

## Introduction

Chemokine receptors belong to the subfamily of class A G protein-coupled receptors (GPCRs), the most therapeutically targeted class of proteins, representing one third of all FDA-approved (Hauser et al., 2017). The majority of chemokines is secreted under inflammatory conditions, and acts as chemoattractant to drive migration of leukocytes to the site of inflammation through chemokine receptors (Scholten et al., 2012). Due to their role in the regulation of the immune system and overexpression in various tumors, chemokine receptors are considered interesting drug targets in a number of therapeutic areas, including e.g. cancer, cardiovascular disorders and multiple sclerosis (Cui et al., 2020, Drouillard et al., 2023, Mikolajczyk et al., 2021, Scholten et al., 2012).

Chemokine receptors typically signal via the G_αi_ signaling pathway (Patel et al., 2013). The atypical chemokine receptor 3 (ACKR3), initially described to as an orphan receptor (Libert et al, 1990), was discovered to act as second high-affinity receptor for CXCL12, next to CXCR4, and subsequently renamed as chemokine receptor CXCR7 (Balabanian et al, 2005). Extensive cell signaling and structural biology studies have now provided evidence that ACKR3 does not signal via the canonical G-protein pathway of chemokine receptors, but only through β-arrestin recruitment (Canals et al., 2012, Rajagopal et al., 2010, Kleist et al., 2022, Yen et al., 2022), and therefore belongs to the family of atypical chemokine receptors (Nibbs and Graham, 2013, Sarma et al., 2023). Next to its β-arrestin-biased signaling, ACKR3 is believed to act as decoy receptor for its endogenous chemokine ligands, CXCL11 (Naumann et al., 2010) and CXCL12 (Balabanian et al., 2005) and various opioid peptides (Meyrath et al., 2020). By sequestrating chemokine receptor ligands, ACKRs are crucial for many physiological processes, such as immune homeostasis and cell migration, as well as for pathological processes such as cancer and inflammation (Burns et al., 2006; Nibbs and Graham, 2013; Ulvmar et al., 2011, Torphy et al., 2022).

With ACKR3 being recognized as interesting drug target (Scholten et al., 2015; Wang et al., 2018, Torphy et al., 2022), the search for different ACKR3 modulators has so far resulted in a variety of modalities that can target ACKR3, including e.g. antibodies and nanobodies, cyclic and opioid peptides and various small-molecule ligands (Adlere et al., 2019, Bobkov et al., 2019, Meyrath et al., 2020). Some of these new modalities bind ACKR3 with high affinity and are offering important chemical biology tools to study ACKR3. Yet, the study of ACKR3 pharmacology still relies extensively on the use of radioactive or fluorescently labelled chemokines (Hatse et al., 2004, Hopkins etl., 2022, Wijtmans et al., 2012, Meyrath et al., 2020). Hence, new approaches to screen for molecules targeting ACKR3 and to study the ACKR3 protein are of interest. Previously, we have reported on a series of styrene-amide ACKR3 agonists, a compound class inspired by Chemocentryx patents (Burns et al., 2005, Melikian et al., 2004) and exemplified by VUF11207 (Fig 1, Wijtmans et al., 2012a). Since VUF11207 is a racemic mixture, we hypothesized that one of its enantiomers would show improved binding to ACKR3. In this study, we report on the synthesis and pharmacological characterization of the new high-affinity enantiomerically pure ACKR3 agonist VUF15485, the most active enantiomer of VUF11207. Importantly, we also describe its labeling with tritium-methylnosylate, affording the first small-molecule radioligand for ACKR3, [^3^H]VUF15485.

**Figure 1.**
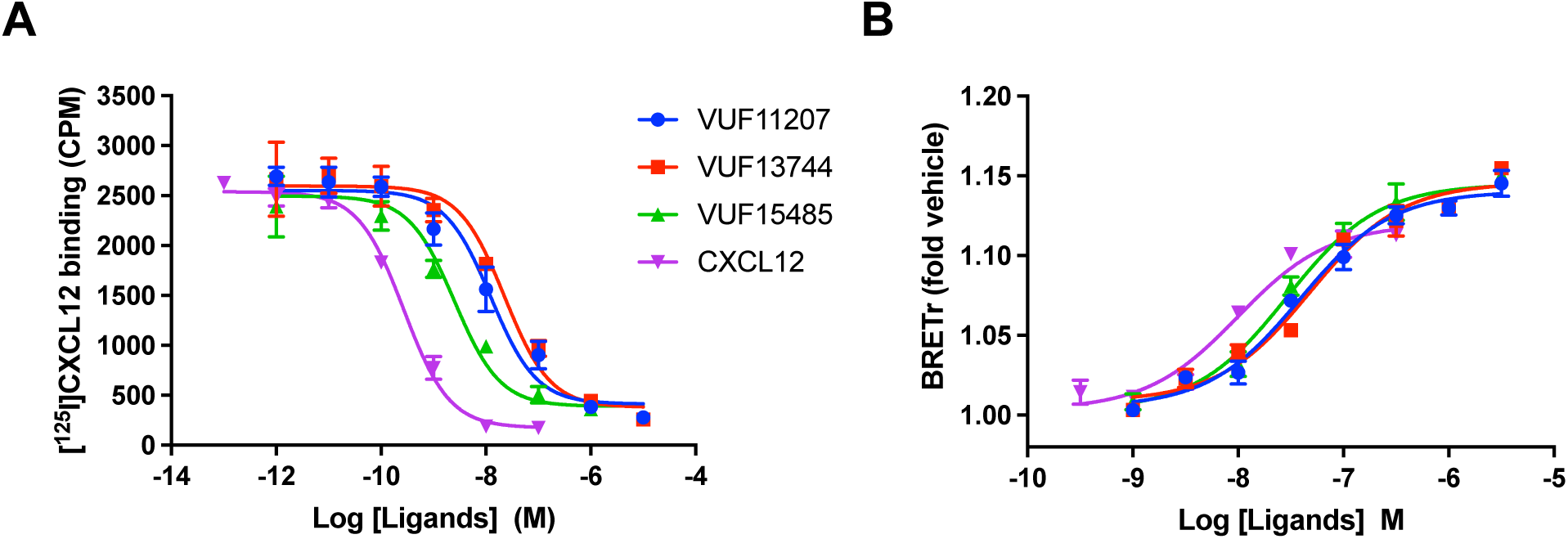
Inhibition of [^125^I]CXCL12 binding and ligand-induced β-arrestin2 recruitment to hACKR3 by CXCL12, VUF11207 and its 2 enantiomers VUF13744 and VUF15485. A) Inhibition of [^125^I]CXCL12 binding by increasing concentrations of unlabeled ligands on intact HEK293T cells expressing hACKR3. B) β-arrestin2 recruitment to hACKR3 in HEK293T cells in response to increasing ligand concentrations as measured by BRET after 20 min of stimulation. Representative curves of at least three independent experiments with triplicates are shown as mean ± SD.

## Materials and Methods

### Synthesis and radiolabelling of VUF15485, ((*R,E*)-N-(3-(2-fluorophenyl)-2-methylallyl)-3,4,5-trimethoxy-N-(2-(1-methylpyrrolidin-2-yl)ethyl)benzamide)

The synthesis and analytical characterization of the two enantiomers of VUF11207 and the radiolabelled high-affinity enantiomer, [^3^H]VUF15485 are described in detail in the Supplementary Data.

### Cell Culture

Human Embryonic Kidney (HEK)293T cells (ATCC) were grown at 37°C and 5% CO_2_ in Dulbecco’s modified Eagle medium (Gibco) supplemented with 10% fetal bovine serum (Bodinco and Gibco), 50 IU/mL penicillin, and 50 mg/mL streptomycin (Gibco). For the experiments with the ACKR3 conformational sensor, HEK293 cells were maintained as adherent cells using Dulbecco’s Modified Eagle Medium (DMEM) supplemented with 4.5 g/L glucose (Thermo Fisher), 10% (v/v) fetal calf serum (Biochrom), 100 U/mL penicillin (Thermo Fisher), 100 µg/mL streptomycin (Thermo Fisher) and 2 mM L-glutamine (Pan Biotech). Cells were split every 2-3 days and kept at 37°C in a humidified atmosphere with 7% CO2.

### [^125^I]CXCL12 Binding Assay

HEK293T cells (2 x 10^6^) were seeded in a 10-cm dish and transfected the next day, as previously described (Adlere et al., 2019). 0.25 µg of pcDEF_3_-hACKR3 and 4.75 µg of empty pcDEF_3_ DNA were combined with 40 µg of 25 kDa linear polyethylenimine (Polysciences) in a total volume of 500 µL 150 mM NaCl and incubated for 20 minutes at room temperature. Subsequently, the DNA/PEI mix was added to the cells upon refreshment of the culture medium. The following day, cells were collected and transferred (120.000 cells per well) into a poly-L-lysine coated (Sigma) 96-well clear plate (Greiner Bio One, PS, F-bottom, clear). On the fourth day, cells were incubated with approximately 50 pM [^125^I]CXCL12 (PerkinElmer) in the absence or presence of unlabeled ligands in binding buffer (50 mM HEPES, pH 7.4, 1 mM CaCl_2_, 5 mM MgCl_2_, 100 mM NaCl, and 1.0% (w/v) bovine serum albumin (BSA, fraction V)) for 3 hours on ice. The incubations were terminated by two rapid washes using ice-cold wash buffer (binding buffer supplemented with 0.5 M NaCl) to separate free from bound radioligand. Next, the cells were lysed using RIPA buffer (0.5 % (v/v) nonidet-P40, 0.5 % (w/v) sodium deoxicholate, 0.1 % (w/v) sodium dodecyl sulfate) and transferred to counting vials (Sarstedt). Recovered radioactivity was determined using a 1282 Compugamma GS (LKB-Wallac).

### [^3^H]VUF15485 Binding Assay

Two days after transfection, hACKR3-expressing HEK293T cells were collected in ice-cold PBS and centrifuged at 1500 g for 10 min at 4°C. The cells were washed with PBS and centrifuged at 1500 g for 10 min at 4°C. Next, membranes were prepared by resuspending the pellet in ice-cold membrane buffer (15 mM Tris pH 7.5, 1 mM EGTA, 0.3 mM EDTA, 2 mM MgCl_2_) followed by homogenization using a teflon-glass homogenizer and rotor (10 strokes at 1100–1200 rpm), as previously described (Adlere et al., 2019). The membranes were subjected to two freeze-thaw cycles using liquid nitrogen and centrifuged at 40,000 g for 25 min at 4°C. The pellet was resuspended in cold Tris-sucrose buffer (20 mM Tris pH 7.4, 250 mM sucrose), and frozen in liquid nitrogen. Protein concentration was determined using a BCA protein assay kit (Thermo Fisher). Membranes expressing hACKR3 (8 ug of total protein) were incubated in 96-well clear plates (Greiner Bio One, PS, U-bottom, clear) with [^3^H]VUF15485 in the absence or presence of unlabeled ligands in binding buffer (10 mM KH_2_PO_4_, 50mM Na_2_HPO_4_, 0.2% (w/v) bovine serum albumin (BSA, fraction V)) at 25°C with gentle agitation. In saturation binding experiments, increasing concentrations of [^3^H]VUF15485 were used, and 10 µM unlabeled VUF15485 was used for determining unspecific binding after 2 hours. In association binding experiments, [^3^H]VUF15485 was incubated for different amounts of time. In dissociation binding, [^3^H]VUF15485 was preincubated for 30 minutes before addition of 10 µM unlabeled VUF15485 and using incubation for different amounts of time. In homologous competition binding experiments, four different concentrations of radioligand were each combined with a concentration series of cold VUF15485 and the mixtures were incubated for 2 hours. In heterologous displacement experiments, 5 nM [^3^H]VUF15485 was combined with a concentration series of cold ligands and incubated for 2 hours. The incubations were terminated by rapid filtration through Unifilter 96-well GF/C plates (PerkinElmer) presoaked with 0.5% PEI using ice-cold wash buffer (50 mM HEPES, pH 7.4, 1 mM CaCl2, 5 mM MgCl2, 500 mM NaCl) to separate free from bound radioligand. The filter plates were dried at 52°C and 25 µL Microscint-O was added. Bound radioactivity was quantified with a MicroBeta scintillation counter (PerkinElmer).

### Bioluminescence Resonance Energy Transfer (BRET)-based β-arrestin2 Recruitment Assay for hACKR3

0.4 µg of pcDEF_3_-hACKR3-RLuc (Canals et al., 2012) and 1.6 µg pcDEF_3_-β-arrestin2-mVenus (Nijmeijer et al., 2013) plasmids were combined to 12 µg of PEI in a total volume of 250 µl 150 mM NaCl and incubated for 20 minutes at room temperature. Thereafter, 1 million resuspended HEK293T cells were added to the DNA/PEI mix, and cells were subsequently seeded (30.000 cells per well) in a poly-L-lysine coated (Sigma) 96-well white plate (Greiner Bio One, PS, F-bottom, white). Two days after transfection, culture medium was substituted with Hanks’ balanced salt solution (Gibco). Next, cells were pre-incubated in Hanks’ balanced salt solution with increasing concentrations of compound for 60 minutes before stimulation with 10 nM CXCL12 and addition of 2 µM Renilla Luciferase substrate coelenterazine-h (Promega). After 20 minutes, RLuc (480/20 nm) and BRET (540/40 nm) signals were measured on the Mithras LB940 (Berthold Technologies). BRET ratios were calculated as BRET/Rluc signal.

### Nanoluciferase complementation-based β-arrestin2 recruitment to chemokine receptors

Comparison of VUF15485-induced β-arrestin2 recruitment to human and mouse ACKR3 was also measured by NanoLuc complementation assay (NanoBiT, Promega Corporation, Madison, WI, USA) in HEK293T cells (Dixon et al., 2016). Briefly, 5 × 10^6^ HEK293T cells were seeded in 10-cm culture dishes and 24 h later co-transfected with pNBe vectors 1 encoding human or mouse ACKR3 C-terminally fused to SmBiT and human or mouse β-arrestin-2 N-terminally fused to LgBiT (Szpakowska et al., 2018). Twenty-four hrs after transfection cells were harvested, incubated for 15 minutes at 37 °C with 200-fold diluted Nano-Glo Live Cell substrate, and distributed into white 96-well plates (5 × 10^4^ cells per well). Cells were then treated with compounds at concentrations ranging from 0.3 nM to 100 µM. Ligand-induced, β-arrestin2 recruitment to receptors was evaluated by measuring bioluminescence with a Mithras LB940 luminometer (Berthold Technologies). For concentration–response curves, the signal recorded was compared to values for human or murine CXCL12 at 300 nM considered as full agonist reference ligand (E_max_ = 100 %). Concentration–response curves were fitted to the three-parameter Hill equation using an iterative, least-squares method (GraphPad Prism version 9.3.1). All curves were fitted to data points generated from the mean of at least three independent experiments.

### Chemokine receptor specificitiy

β-arrestin2 recruitment to human chemokine receptors in response to 1 µM VUF15485 or 200 nM positive control chemokines was also monitored by NanoLuc complementation assay (NanoBiT, Promega) (Dixon et al., 2016), as previously described (Szpakowska et al., 2018). In brief, 6,5 × 10^5^ HEK293T cells were plated per well of a 12-well dish and 24 hours later co-transfected with pNBe vectors encoding a human chemokine receptor C-terminally tagged to SmBiT and human β-arrestin2 N-terminally fused to LgBiT (Meyrath et al., 2020). This was adjusted in case of the CXCR4, in which the β-arrestin2 was N-terminally fused to the SmBiT and CXCR4 was C-terminally fused to the LgBiT. Cells were harvested 24 hours after transfection, incubated 25 min at 37 °C with Nano-Glo Live Cell substrate diluted 200-fold and distributed into white 96-well plates (1 × 10^5^ cells per well). Ligand-induced, human β-arrestin2 recruitment to chemokine receptors was monitored with a Mithras LB940 luminometer (Berthold Technologies) for 20 min. For each receptor, 200 nM of one known agonist chemokine listed in the IUPHAR repository of chemokine receptor ligands was added as positive control (see details in Meyrath et al., 2020). To evaluate the antagonist properties of VUF15485, full agonists of each receptor (20 nM except for CCR9, CCR10, XCR1 (each 200 nM) were added after the 20 min incubation with VUF15485. Signal from wells treated with full agonist only was defined as 100 % activity and signals from wells treated with no agonist were used to set 0% activity.

### Confocal Microscopy

To investigate the expression and localization of the ACKR3 conformational sensor, HEK293 cells were seeded on 24 mm glass cover slips placed in wells of a 6-well plate. The coverslips had been previously coated with poly-D-lysine (1 mg/ml) for 30 minutes and washed once with PBS. The day after, cells were transfected with 0.3 µg HA-ACKR3-FlAsH-CFP and, when indicated, 0.6 µg K44A dynamin using Effectene (Qiagen) and following the manufacturer’s instructions. After 24h, cells were analyzed on a confocal microscope while kept in 600 µL of measuring buffer (140 mM NaCl, 5.4 mM KCl, 2 mM CaCl_2_, 1 mM MgCl_2_, 10 mM HEPES; pH 7.3). Confocal microscopy was performed using a Leica TCS SP8 system with an Attofluor holder (Molecular Probes) and images were taken with a 63× water objective (numerical aperture, 1.4). The CFP was excited with a diode laser line at 442 nm and the emission fluorescence intensity was recorded from 450 to 600 nm. Images were acquired at 1024 ×1024 pixel format, line average 3, and 400 Hz.

### FlAsH labeling

FlAsH labeling was performed as previously described (Hoffmann et al., 2010). Briefly, transfected cells grown on 40 mm Willco dishes were washed twice with labeling buffer (150 mM NaCl, 10 mM HEPES, 2.5 mM KCl, 4 mM CaCl2, 2 mM MgCl2, freshly supplemented with 1 g/L glucose; pH 7.3). After, cells were incubated for 1 h at 37°C with labeling buffer containing 1 µM FlAsH and 12.5 µM 1,2-ethanedithiol (EDT). This step allows FlAsH to bind to its binding site in the sensor. Then, cells were rinsed twice and incubated for 10 min at 37°C in labeling buffer containing 250 µM EDT to reduce non-specific FlAsH labeling. Finally, cells were washed twice again and kept in media at 37°C until measurement.

### Determination of the FRET efficiency and dynamic FRET measurements

The FRET efficiency was investigated as described (Jost et al., 2008). To study the intramolecular FRET in the HA-ACKR3-FlAsH-CFP sensor, HEK293 cells were prepared in coverslips as described above for confocal microscopy and co-transfected with 0.3 µg HA-ACKR3-FlAsH-CFP and 0.6 µg K44A dynamin per dish. After 24h, cells were FlAsH-labeled as described. Next, the coverslips were mounted on the Attofluor holder and 999 µL of measuring buffer were added. After recording for approximately 40 s, 1 µL of British Anti-Lewisite (BAL; Sigma) was added to the buffer solution to reach a final concentration of 5 mM. This concentration of BAL displaced FlAsH from its binding site in the sensor and dequenced the CFP. FRET efficiency was then calculated by analyzing the increase detected in the CFP emission using the formula ((FCFPmax/(FCFPmin-FCFPmax)). To study the intermolecular FRET of the sensor, HEK293 cells were co-transfected with 0.2 µg HA-ACKR3-FlAsH, 0.2 µg HA-ACKR3-CFP and 0.8 µg of K44A dynamin, and the same procedure was followed. Data was analyzed using the Clampfit software (version 10.7; Molecular Devices).

For measuring activation kinetics using the receptor sensor, HEK293 cells were seeded on 40 mm Willco dishes that had been previously coated with poly-D-lysine for 30 minutes and washed with PBS once. After 24 h, cells were co-transfected with 0.6 µg HA-ACKR3-FlAsH-CFP and 1.2 µg K44A dynamin per dish using the Effectene reagent (Qiagen) and following the manufacturer’s instructions. 24h after transfection, cells were FlAsH-labeled. During the experiment, cells were kept in measuring buffer and superfused with buffer or buffer supplemented with the appropriate ligand (CXCL12 or VUF15485) at the indicated concentration using the BioPen® microfluidic system from Fluicell (Ainla et al., 2010; Ainla et al., 2012, Perpina-Viciano et al., 2020). The tau values (apparent on-rate) were determined using the Clampfit by fitting a mono-exponential decay function to the FRET change detected upon ligand stimulation. Data was corrected for direct excitation of the acceptor, donor bleed-through and photobleaching using OriginPro (OriginLab). FRET measurements were performed using the microscopic FRET set-up as described (Hoffmann et al., 2005).

### Data analysis

Data was analyzed by non-linear regression using GraphPad Prism 7 software. Association binding curves were fitted using the “association kinetics – two or more conc. of hot” equation, and dissociation binding were fitted using the “Dissociation – one phase exponential decay” equation. Saturation binding curves were fitted using the “one site – total and nonspecific binding” equation. Competition binding curves were fitted using the “one site – fit logIC50” equation. BRET concentration response curves were fitted using the “log(agonist) vs. response (three parameters)” equation. All values shown are mean ± SD values of at least 3 independent experiments.

### ACKR3 site-directed mutagenesis

For the site-directed mutagenesis studies the DNA sequences of the wild-type ACKR3 (NM_020311.3), N-terminally fused to an HA-tag in the expression vector pcDEF3 was used as template. Residue numbering is displayed throughout the manuscript as absolute UniProt sequence numbers and with the Ballesteros–Weinstein notation (Ballesteros and Weinstein, 1995) in superscript, in which the first number denotes the helix, the second the residue position relative to the most-conserved residue, defined as number 50, in a non-gapped sequence alignment. The single amino acid ACKR3 mutants Y51^1.39^A, W100^2.60^Q, S103^2.63^D, H121^3.29^A, F124^3.32^A, D179^4.60^N, E213^5.39^Q, V217^5.43^, Y268^6.51^A, D275^6.58^N, H298^7.36^A, Q301^7.39^E were generated by PCR-based mutagenesis and the sequences were verified by DNA sequencing. Expression of mutant receptors at the cell membrane was verified by radioligand binding studies and anti-HA ELISA as described previously (Verweij et al., 2020).

### VUF15485 docking

The ACKR3 protein was prepared in Internal Coordinate Mechanics (ICM) version 3.9-3a (Abagayan et al., 1994) using the cryo-EM structure of ACKR3 in complex with the partial agonist CCX662 and bound to an intracellular Fab (PDB entry 7SK9, 3.8 Å resolution) as template (Yen et al., 2022). The Fabs CID25, CID24 and cholesterol molecules present in the structure were deleted. The residues in the vicinity (5.0 Å) of the small molecule CCX662 were optimized for the hydrogen bond network and minimized (cartesian minimization in Merck molecular force field, MMFF). VUF15485 was docked and scored using the ICM VLS scoring function (Neves et al., 2012).

## Results

### Pharmacological evaluation of VUF11207 enantiomers

We previously reported VUF11207 to act as a potent small-molecule ACKR3 agonist (Wijtmans et al., 2012a). However, VUF11207 is racemic and for the development of a suitable radioligand the isolation and pharmacological characterization of its two enantiomers was required. Enantiopure isomers of VUF11207, i.e. VUF13744 and VUF15485, (Scheme 1) were prepared based on our previously disclosed synthesis route (Wijtmans et al., 2012a) by starting with the respective enantiopure amine precursors (see Supplementary Data). First, we determined the affinity for hACKR3 of the two enantiomers VUF13744 and VUF15485 using a [^125^I]CXCL12 displacement assay on intact HEK293T cells, transiently transfected with pcDEF3-hACKR3 (Wijtmans et al., 2012a). VUF13744 and VUF15485 both displace the binding of labelled chemokine to hACKR3 almost to the same level as unlabeled CXCL12, resulting in pIC_50_ values of 7.7 ± 0.1 and 8.3 ± 0.1, respectively (Fig 1A, pIC_50_ of CXCL12 = 9.6 ± 0.1). As expected, racemate VUF11207 displaces [^125^I]CXCL12 binding to hACKR3 with a pIC_50_ value in between those of its two enantiomers (pIC_50_ = 8.0 ± 0.1, Fig 1A).

**Scheme 1.**
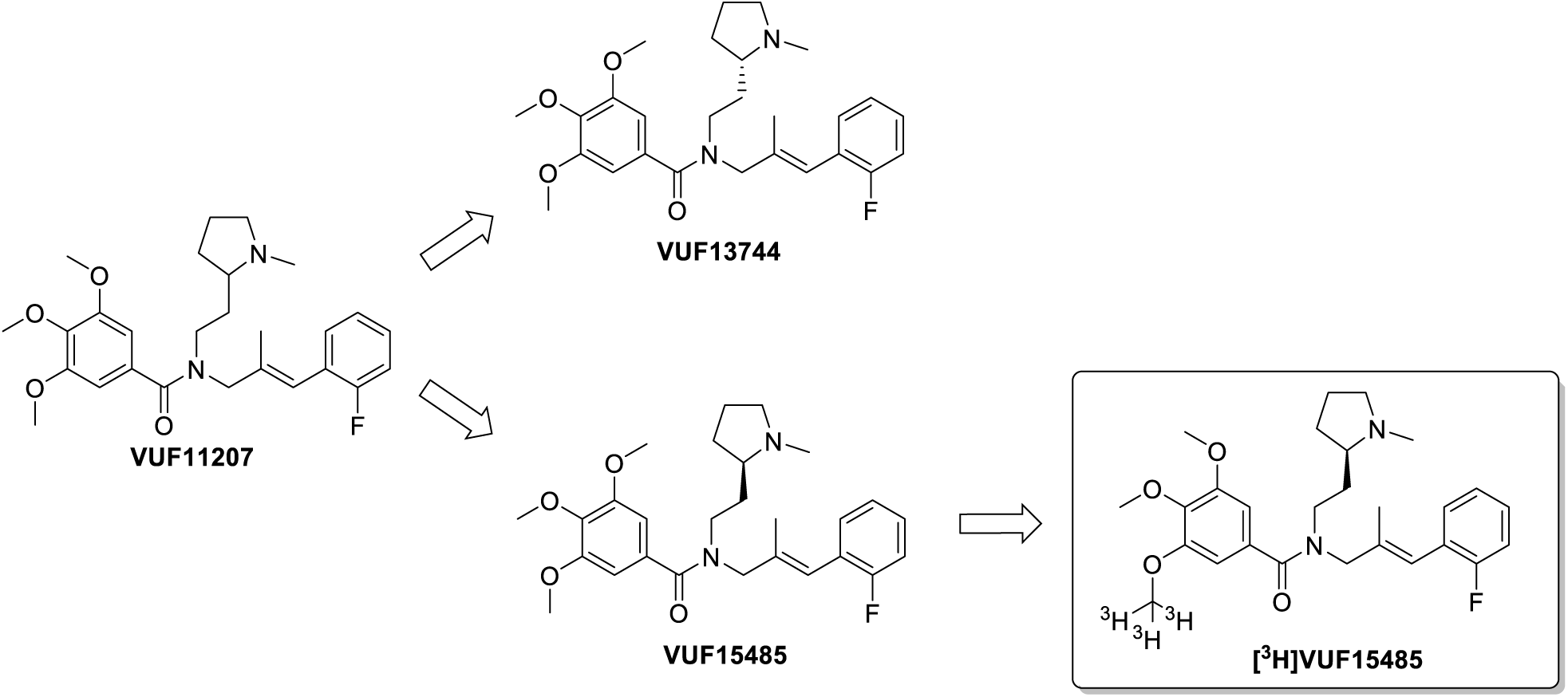
Structures of VUF11207, its two stereoisomers and the radiolabeled [^3^H]VUF15485

Next, the potency of the three compounds was determined using a BRET-based β-arrestin2 recruitment assay. As previously reported (Wijtmans et al., 2012a), both CXCL12 and VUF11207 effectively recruit β-arrestin2 to hACKR3 with pEC_50_ values of 8.2 ± 0.0 and 7.5 ± 0, respectively (Fig 1B). In this functional assay, VUF15485 has a slightly higher potency than VUF13744 with pEC_50_ values of 7.6 ± 0.1 and 7.4 ± 0.1, respectively (Fig. 1B). After 20 minutes of stimulation, CXCL12 (pEC_50_ = 8.2 ± 0.0) exhibits a slightly lower efficacy (Fig. 1B) compared to the three compounds. Moreover, CXCL12 and VUF15485 also effectively recruited β-arrestin2 to mouse ACKR3 with pEC_50_ values of 8.7 ± 0.1 and 7.7 ± 0.1, respectively. At the mACKR3 VUF15485 recruited β-arrestin2 slightly less efficacious (α = 0.8 ± 0.1, n = 3).

To study ACKR3 activation upon ligand binding with high temporal resolution, we developed a FRET-based sensor for hACKR3 as shown in figure 2A. We used the cyan fluorescent protein (CFP) in combination with fluorescein arsenical hairpin binder (FlAsH), a small cell-permeant fluorescein-derivative. This combination of fluorophores has been successfully used before to investigate receptor dynamics in living cells and evaluate ligand binding and ligand efficacy in real time for other GPCRs (Stumpf and Hoffmann, 2016). In the current study, the CFP was fused to the C-terminal end of HA-tagged hACKR3 and the FlAsH-binding sequence **CCPGCC** was introduced in the predicted third intracellular loop (ICL-3) between residues Ser^242^ and Ser^243^. Subsequently, we assessed the affinity of the ACKR3 sensor for CXCL12, its expression, localization and FRET efficiency of the ACKR3 sensor. After transient expression in HEK293 cells, CXCL12 exhibits similar affinity for wild type (WT) receptor (pIC_50_ = 9.0 ± 0.2) and the hACKR3 sensor (pIC_50_ = 9.1 ± 0.2), as determined by means of [^125^I]CXCL12 displacement on membranes (sup. fig. S1B). These data indicate that ligand binding to the sensor is not affected by the presence of the FlAsH and CFP fluorophores on the intracellular side, which is in agreement with previous studies (Hoffmann et al., 2005). The hACKR3 has been shown to localize predominantly to intracellular compartments of different cell types (Canals et al., 2012, Luker et al., 2010; Shimizu et al., 2011, Zarca et al., 2021). As expected, confocal images of HEK293 cells expressing the HA-ACKR3-FlAsH-CFP construct showed that the sensor is predominantly found in intracellular compartments in the cytoplasm of these cells (supp. fig. S1C). Therefore, we co-transfected the sensor with K44A dynamin to block GPCR endocytosis (Ray et al., 2012) and to make it accessible for ligands. Indeed, this approach significantly increased the localization of the HA-ACKR3-FlAsH-CFP sensor on the plasma membrane (supp. fig. S1C). As VUF15485 showed the highest affinity, it was characterized along with CXCL12 using this newly generated conformational sensor. In line with the similar agonistic profiles of both agonists in the β-arrestin2 recruitment assay (vide supra), VUF15485 triggers similar conformational changes as CXCL12 (Fig. 2C-D), although with different tau values. The observed conformational change of ACKR3 upon activation by VUF15485 exhibits slower kinetics than after activation with the chemokine CXCL12 (Fig. 2B).

**Figure 2.**
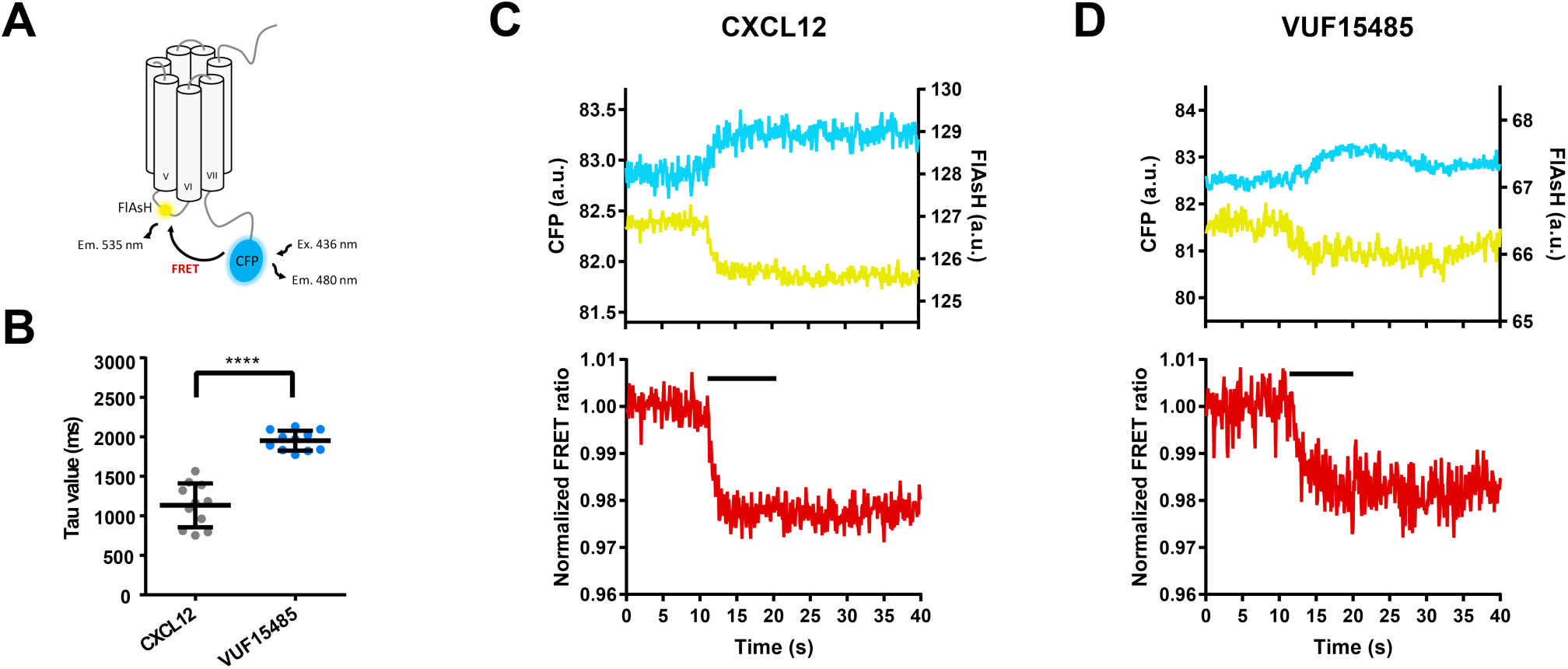
Kinetic analysis of ACKR3 activation in response to CXCL12 and VUF15485. A) Schematic depiction of the hACKR3-based FRET sensor. Upon excitation of the CFP at 436 nm, energy is efficiently transferred to FlAsH, resulting in FRET. Upon ligand binding, the receptor undergoes a conformational change, simultaneously altering the distance and/or orientation between the fluorophores, which ultimately leads to a change in FRET. B) On-kinetics of ACKR3 activation in response to the different ligands. Tau values were determined from the normalized FRET ratios of individual experiments (e.g. panel C and D) by fitting the change detected to a mono-exponential decay function, and represented in a scatter plot. Data shows the mean ± SD of 11 cells, measured on at least three independent experimental days. C-D) Traces measured from a single HEK293 cell expressing the ACKR3-FlAsH-CFP sensor and K44A dynamin and stimulated with 30 µM CXCL12 (C) or 50 µM VUF15485 (D). The black short lines indicate stimulation with the ligand. The presented data is representative. Upper panels show the FlAsH (yellow color) and CFP (cyan color) emissions recorded at 480 and 535 nm, respectively. Lower panels show the corrected and normalized FRET ratio in red.

### Pharmacological specificity of VUF15485

For the use of VUF15485 as a selective agonist of the ACKR3, it is important to know whether its activity is specific for ACKR3. Using β-arrestin2 recruitment as a functional readout of receptor activity, the effect of 1 µM VUF15485 on receptor activity was investigated for the complete panel of chemokine receptors, transiently expressed in HEK293T cells (Fig. 3). To do so, chemokine receptors and β-arrestin2 were fused to complementary fragments of the NanoLuc protein which can reconstitute into a functional luminescent enzyme when in close proximity (Meyrath et al., 2020). Upon treatment of cells with 1 µM VUF15485, clear β-arrestin2 recruitment to the ACKR3 was measured, but not to any of the other chemokine receptors, indicating high specificity of agonism (Fig. 3A). Moreover, antagonism of the activated chemokine receptors by 1 µM VUF15485 was in all cases less than 50% (Fig. 3B). Hence, at saturating concentrations (>60x *K*_d_ at the ACKR3) VUF15485 has very limited effect on chemokine receptors other than ACKR3.

**Figure 3.**
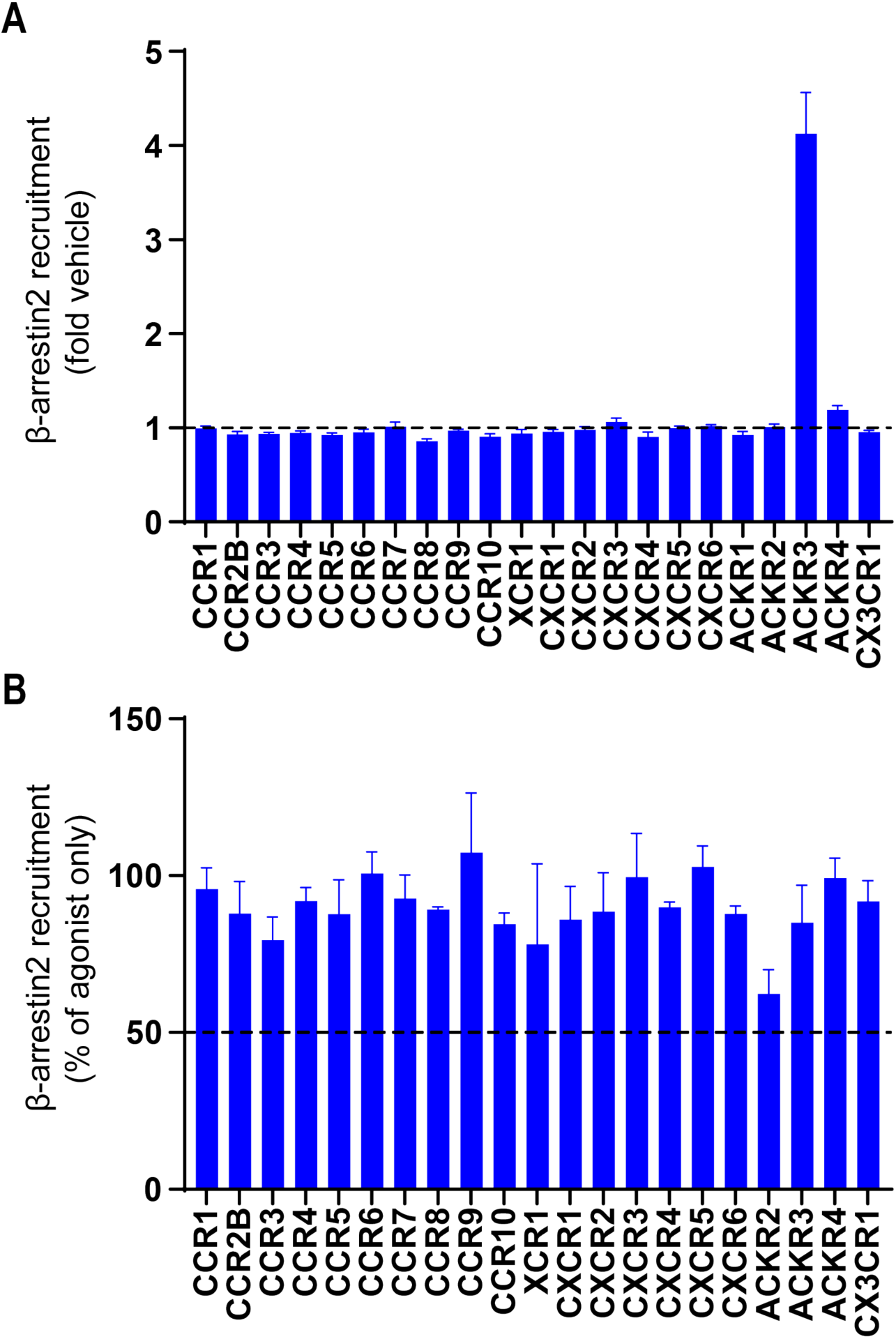
VUF15485 specifically modulates the ACKR3 over other chemokine receptors. HEK293T cells were transfected with β-arrestin2 together with one of the chemokine receptors, each fused to one of the components of the Nanobit nanoluciferase complementation sensors. (A) Chemokine receptor activation by 1 µM VUF15485 was measured as the recruitment of β-arrestin2 to the indicated chemokine receptor. Data show the fold increase in luminescence over vehicle. (B) The relative inhibition of the endogenous agonist induced β-arrestin2 recruitment was determined for each chemokine receptor upon treatment with 1 µM VUF15485. Data show the signal of VUF15485 treated cells after endogenous agonist treatment normalized in % to the signal from endogenous agonist treatment alone. The graphs represent mean and SEM of ≥ 3 experiments.

### Radiolabeling of VUF15485 with tritium

Since VUF15485 was identified as the most potent enantiomer of VUF11207 at hACKR3, VUF15485 was selected for radiolabeling with tritium. The structure of VUF15485 provides multiple methyl groups that could be considered for late-stage CT_3_ incorporation. However, installing a CT_3_ group on the amine was discarded due to the very limited availability of suitable enantiopure secondary amine precursors. Rather, introducing a CT_3_ unit on one of two possible phenol precursors was deemed a better approach. After selection of the optimal phenol group, methylation was conducted using [^3^H]methylnosylate affording [^3^H]VUF15485 (Scheme S1, Supplementary Data) resulting in 38 MBq (38% radiochemical yield) with a radiochemical purity of 98% and a molar activity of 2.98 MBq/nmol, based on the molar activity of ^3^H]methylnosylate.

### Assay development for [^3^H]VUF15485 binding to hACKR3

First, we optimized the binding conditions for [^3^H]VUF15485 using membranes from pcDEF3-hACKR3 transfected HEK293T cells. Different buffer compositions (buffer 1: 10 mM KH_2_PO_4_, 50 mM Na_2_HPO_4_; buffer 2: 50 mM HEPES, 1 mM CaCl_2_, 5 mM MgCl_2_, 100 mM NaCl, pH 7.4; buffer 3: 50 mM Tris-HCl, pH 7.4, 100 mM NaCl, 0.1% (w/v) Tween80) were initially tested to maximize total and minimize non-specific binding (determined by addition of [^3^H]VUF15485 in the absence or presence of a saturating concentration of 10 µM unlabeled VUF15485). Buffer 1 (10 mM KH_2_PO_4_, 50 mM Na_2_HPO_4_) gave the best results (data not shown) and was used thereafter for optimization of the binding assay. To increase specific binding of [^3^H]VUF15485, addition of bovine serum albumin (BSA) or Tween 20 was evaluated (Fig 4A). The latter almost completely abolished specific binding of the radioligand. Addition of BSA did not increase the specific binding dramatically (fig. 4A), but may in general prevent non-specific binding of compounds and proteins by adsorption to plastic (Flanagan, 2016) and was therefore maintained. Next, the binding of [^3^H]VUF15485 to increasing amounts of hACKR3 expressing membranes was measured under these conditions (Fig. 4B). The specific binding of the radiolabel linearly increased with increasing amounts of membranes.

**Figure 4.**
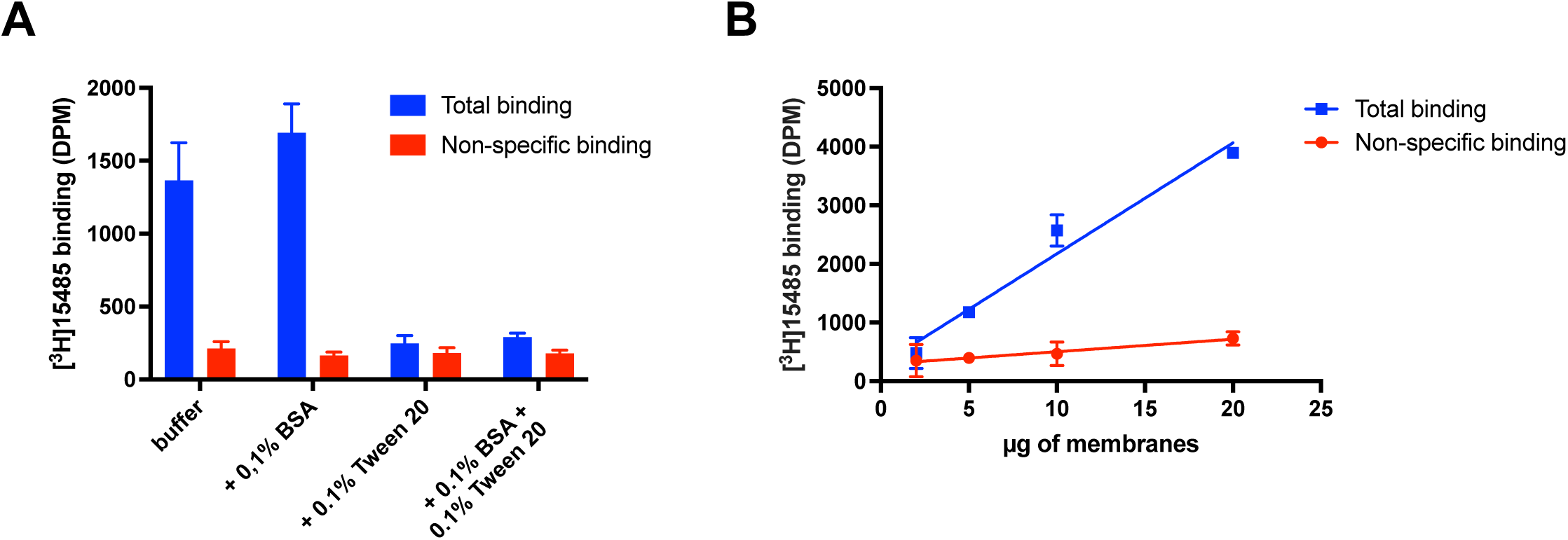
Optimization of assay conditions for [^3^H]VUF15485 binding to hACKR3. A) Optimization of binding buffer (50 mM Na_2_HPO_4_, 10 mM KH_2_PO_4_) by supplementation with 0.1% BSA and/or 0.1% Tween 20 to increase the specific binding. Total and non-specific (determined in the presence of 10 µM of VUF15485) binding of 5 nM [^3^H]VUF15485 to 8 µg hACKR3-expressing HEK293T membranes. B) Total and non-specific binding (determined in the presence of 10 µM of VUF15485) of 5 nM of [^3^H]VUF15485 to increasing amount of hACKR3-expressing membranes in optimized binding buffer (50 mM Na_2_HPO_4_, 10 mM KH_2_PO_4_, 0.1% BSA). Representative curves of 3 independent experiments, each performed in triplicates plotted as mean ± SD are shown.

### Characterization of [^3^H]VUF15485 binding to hACKR3

After assay optimization, we determined the pharmacological properties of the radioligand. To determine the equilibrium affinity constant (*K*_d_) for [^3^H]VUF15485 binding to hACKR3, association and dissociation kinetic experiments were performed to obtain the on-rate (*k*_on_), the off-rate (*k*_off_) (Fig. 5A, 5B), and the calculated kinetically derived *K*_d_ values (defined as *K*_d_ = *k*_off_/*k*_on_). To measure its on-rate, the radioligand was incubated with membranes expressing hACKR3 until equilibrium was reached (Fig. 5A). [^3^H]VUF15485 proved to associate relatively fast to hACKR3 with equilibrium being reached after only 4 minutes using 5 nM of radioligand, providing a *k*_on_ value of 0.092 ± 0.008 min^-1^ nM^-1^ (n = 3). To determine the off-rate of the radioligand, [^3^H]VUF15485 was incubated with membranes expressing hACKR3 until an equilibrium was reached, and dissociation of the compound-receptor complex was induced using a saturating (10 µM) concentration of unlabeled VUF15485 (Fig. 5B). Similar to the association kinetics, [^3^H]VUF15485 dissociates rapidly from the receptor, with complete dissociation of the compound after five minutes, giving a *k*_off_ value of 0.754 ± 0.03 min^-1^ (n = 3). The resulting target residence time (RT) of [^3^H]VUF15485, defined as 1/*k*_off_, amounts to only 1.32 minutes. Based on the fitted on- and off-rate constants, the *K*_d_ value for the binding of [^3^H]VUF15485 to hACKR3 was calculated to be 8.2 ± 0.8 nM.

**Figure 5.**
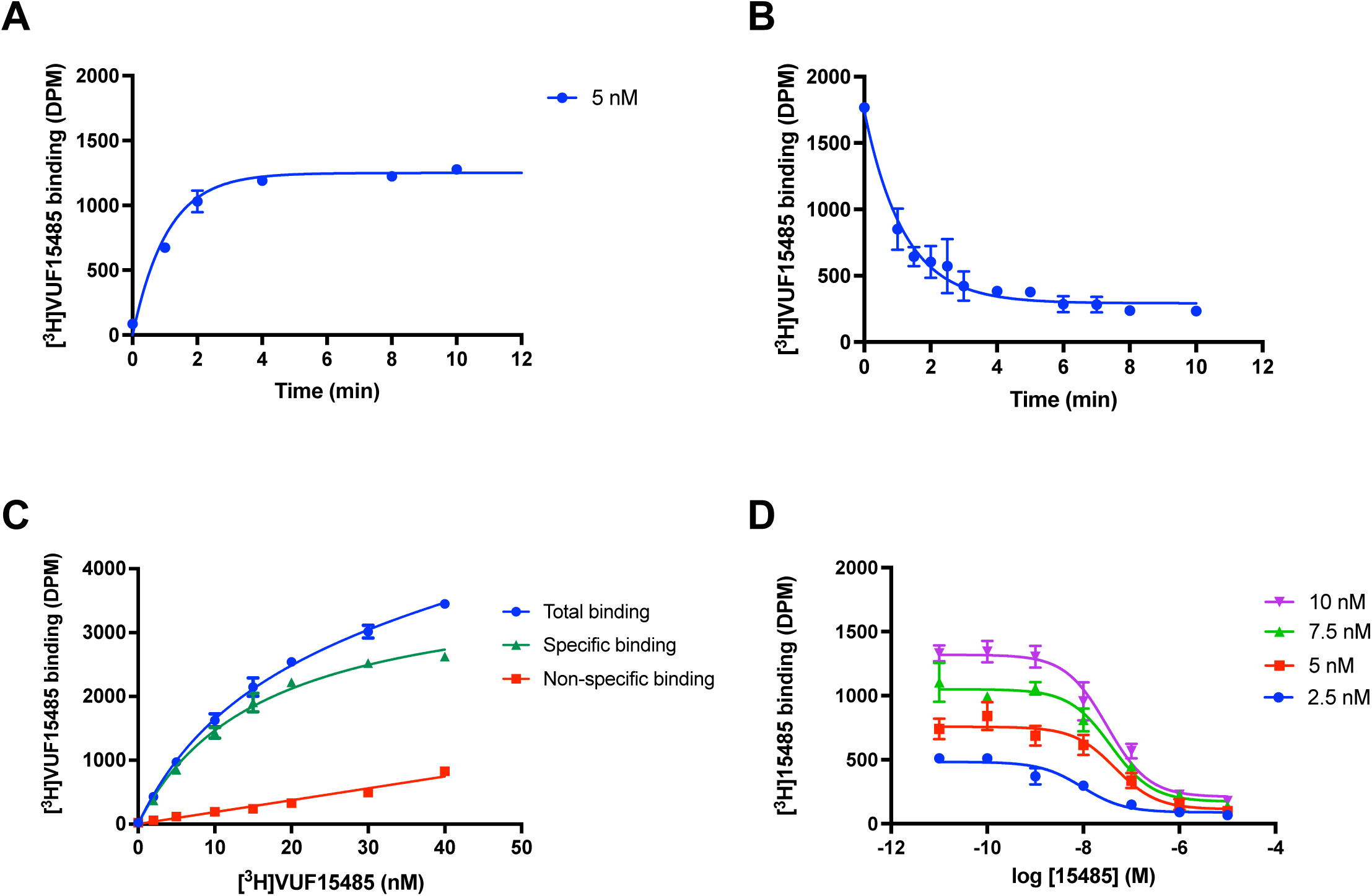
Pharmacological characterization of [^3^H]VUF15485 binding to hACKR3. A) Association binding kinetics of 5 nM [^3^H]VUF15485 to hACKR3. B) Dissociation kinetics of pre-bound [^3^H]VUF15485 (7 nM) from hACKR3 were initiated by the addition of 10 µM unlabeled VUF15485 and measured over time. C) Saturation binding of increasing concentrations [^3^H]VUF15485 to hACKR3-expressing membranes. Specific binding was calculated by subtracting non-specific binding in the presence of 10 µM VUF15485 from total binding. D) Homologous competition binding experiments of four concentrations [^3^H]VUF15485 with increasing concentrations of unlabeled VUF15485. Representative curves of at least three independent experiments with triplicates are shown as mean ± SD.

Saturation binding experiment, in which increasing concentrations of [^3^H]VUF15485 were incubated with membranes expressing hACKR3 until equilibrium was reached (Fig. 5C) in the presence or absence of 10 µM of VUF15845 (to determine unspecific binding), revealed a *B*_max_ value of 5.43 ± 1.61 pmol/mg of protein (n = 3) and a *K*_d_ value of 16.0 ± 2.4 nM (n = 3). The latter is slightly higher compared to the kinetically-derived *K*_d_ value.

Next, competition binding experiments were performed to determine the affinity of unlabeled VUF15485 using [^3^H]VUF15485. We performed homologous competition binding experiments using different concentrations of [^3^H]VUF15485 (2.5, 5, 7.5, and 10 nM) in the range of its *K*_d_ value (Fig. 5D). Dilution series of VUF15485 were incubated with membranes expressing hACKR3 in the presence of different concentrations of [^3^H]VUF15485, from which a *K*_i_ value of 33.8 ± 8.6 nM for unlabeled VUF15485 was extrapolated (n = 3).

### Use of [^3^H]VUF15485 to probe the binding of chemokine receptor antagonists to hACKR3

To assess the pharmacological profile of the [^3^H]VUF15485 binding to hACKR3, we selected a panel of small molecules of varying chemotypes (SI Fig S3), that are known to act on hACKR3 and on the related chemokine receptors CXCR3 (also binding CXCL11) and CXCR4 (also binding CXCL12) (Table 1). The panel of chemokine receptor ligands, used in the [^3^H]VUF15485 competition binding experiments (Fig. 6A – 6B), includes 3 small-molecule ACKR3 agonists VUF15485, VUF11403 and VUF16545 (Table 1) (Uto-Konomi et al., 2013, Wijtmans et al., 2012a), CXCR4 antagonists VUF16771, VUF16774, VUF16779 (Adlere et al., 2019), IT1t (Thoma et al., 2008) and the FDA-approved Plerixafor/AMD3100 (Schols et al., 1997), and the CXCR3 antagonists VUF11418, VUF11222, VUF10990 and VUF11211 (Scholten et al., 2014; Scholten et al., 2015; Shao et al., 2011; Wijtmans et al., 2012b), next to the endogenous ACKR3 ligands CXCL11 (Burns et al., 2006) and CXCL12 (Balabanian et al., 2005).

**Figure 6.**
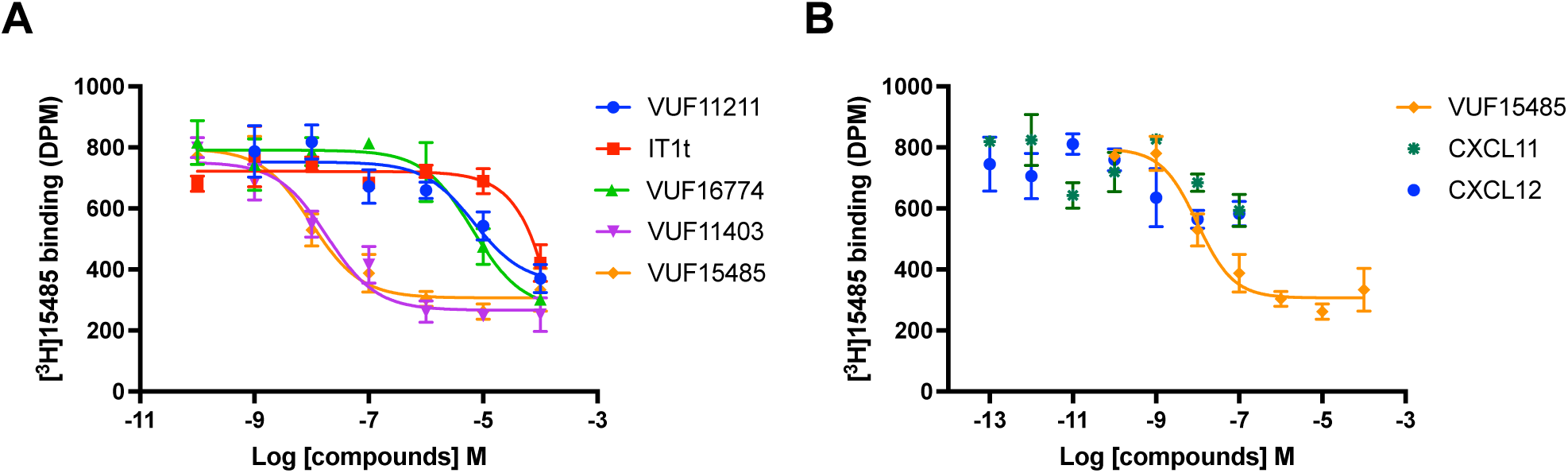
Inhibition of [^3^H]VUF15485 binding to hACKR3 by increasing concentrations of selected unlabeled CXCR3, CXCR4 and ACKR3 chemokine receptor ligands. Representative curves of at least three independent experiments with triplicates are shown as mean ± SD

**Table 1.** Inhibition of [^3^H]VUF15485 binding to hACKR3 by selected unlabeled CXCR3, CXCR4 and ACKR3 chemokine receptor ligands. All results are average ± S.E.M. of at least three independent experiments. The chemical structures of all compounds are displayed in the SI.

|  | **Displacing**<br>**ligand** | **[<sup>3</sup>H]VUF15485 binding**<br>**pIC<sub>50</sub> ± SEM (n ≥ 3)** | **Cmpd code & reference** |
| --- | --- | --- | --- |
| ACKR3<br>ligands | VUF15485 | 7.8 ± 0.1 | this paper |
| | VUF11403 | 7.9 ± 0.1 | **30** in Wijtmans et al., 2012a |
| | VUF16545 | 7.8 ± 0.1 | **1** in Uto-Konimi et al., 2013 |
| CXCR4<br>ligands | VUF16771 | 5.8 ± 0.1 | **28** in Adlere et al., 2019 |
| | VUF16774 | 5.0 ± 0.1 | **9** in Adlere et al., 2019 |
| | VUF16779 | 6.0 ± 0.1 | **22** in Adlere et al., 2019 |
| | IT1t | $\leq 4$ | |
| | AMD3100 | $\leq 4$ | |
| CXCR3<br>ligands | VUF11418 | 5.9 ± 0.3 | **39** in Wijtmans et al., 2012b |
| | VUF11222 | 5.8 ± 0.2 | **38** in Wijtmans et al., 2012b |
| | VUF10990 | 5.9 ± 0.1 | **10q** in Wijtmans et al., 2011 |
| | VUF11211 | 4.7 ± 0.2 | Scholten et al., 2014 |

The affinity values of unlabeled VUF15485 for binding to hACKR3 are comparable when measured by either competition of [^3^H]VUF15485 or [^125^I]CXCL12 (pIC_50_ = 7.8 ± 0.1 and 8.3 ± 0.1 respectively). The same holds true for the affinity for hACKR3 of VUF11403, a compound of the same chemotype as VUF15485 (SI Fig S3), measured by displacing [^3^H]VUF15485 binding (pIC_50_ = 7.9 ± 0.1). This value is also comparable to the previously published data from competition binding experiments using [^125^I]CXCL12 binding to hACKR3 (p*K*_i_ = 7.7 ± 0.1) (Wijtmans et al., 2012a). The binding of [^3^H]VUF15485 to hACKR3 is also effectively displaced by the structurally different (SI Fig S3) ACKR3 agonist VUF16545 (Table 1). Interestingly, the ACKR3 cognate ligands CXCL11 and CXCL12 marginally displace [^3^H]VUF15485 binding to hACKR3 expressing membranes (Fig. 6B). This is in contrast with VUF15485 fully displacing [^125^I]CXCL12 binding to HEK293T cells expressing hACKR3 (Fig. 1A). [^3^H]VUF15485 binding experiments were subsequently used to determine whether previously published CXCR3 and CXCR4 small-molecule ligands show affinity for ACKR3. All tested CXCR3 antagonists were able to displace [^3^H]VUF15485 binding to hACKR3 albeit at high concentrations, indicating they exhibit some (low to high micromolar) affinity for hACKR3. The CXCR3 antagonists VUF10990, VUF11222 and VUF11418, all three from the same chemical class, show pIC_50_ values around 5.8-5.9. On the other hand, VUF11211, a CXCR3 antagonist from a different chemotype, shows a lower affinity for hACKR3 (pIC_50_ = 4.7 ± 0.2). The previously published CXCR4 ligands IT1t and AMD3100 also displace [^3^H]VUF15485 binding to hACKR3 only at high concentrations (Fig. 6A, Table 1). A new series of CXCR4 antagonists recently published by our lab was also tested and shows only (moderate) affinity for hACKR3 (pIC_50_ = 5.0-6.0).

### Site-directed mutagenesis studies and VUF15485 docking

Next, the ACKR3 binding site for the agonist VUF15485 was explored. Initially negatively charged residues in the binding site were mutated to their correspondent non-charged residues: D179^4.60^N, E213^5.39^Q, and D275^6.58^N. Additionally, polar residues S103^2.63^ and Q301^7.39^, involved in ligand binding at chemokine receptors CXCR3 and CXCR4 were mutated to negatively charged residues (D and E respectively). These mutations mimic the closely related CXCR3 and CXCR4, both sharing chemokine ligands with ACKR3 (Scholten et al., 2012). The mutant ACKR3 proteins are all detectable by ELISA on membranes of transiently transfected HEK293T cells (Fig. S4). In homologous [^3^H]VUF15485 displacement studies, VUF15485 binding to hACKR3 results in a pK_i_ value of 7.6 ± 0.07 (Table 2). The high-affinity binding of VUF15845 was not affected by the E213^5.39^Q, D275^6.58^N, S103^2.63^D and Q301^7.39^E mutations. These mutations did also not reduce the affinity of [^125^I]CXCL12 (Table 2). In contrast, [^3^H]VUF15485 binding was completely abolished by the mutation D179^4.60^N in TM4 (fig. 4.5). This negatively charged residue has previously been shown to be involved in the binding of CXCL12 to ACKR3 (Benredjem et al., 2017), and also in this study the D179^4.60^N ACKR3 protein did not bind [^125^I]CXCL12 (Table 2). This mutant is normally expressed on the cell membrane as measured by ELISA (Fig. S4), suggestion that the loss of [^3^H]VUF15485 binding is due to an ionic interaction between VUF15485 and D179^4.60^.

**Table 2.** [^3^H]VUF15485 and [^125^I]CXCL12 binding to selected hACKR3 mutant proteins, transiently expressed in HEK293T cells. The pK_i_ values were determined by homologous competition binding experiments. All results are average ± S.E.M. of three independent experiments.

| Mutant | TM domain | **[<sup>3</sup>H]VUF15485** |  | **[<sup>125</sup>I]CXCL12** |  |
| --- | --- | --- | --- | --- | --- |
|  |  | pK<sub>i</sub> | SEM | pK<sub>i</sub> | SEM |
| WT | - | 7.61 | 0.07 | 8.86 | 0.12 |
| S103<sup>2.63</sup>D | TM2 | 7.02 | 0.13 | 8.89 | 0.17 |
| D179<sup>4.60</sup>N | TM4 | no binding | - | no binding | - |
| E213<sup>5.39</sup>Q | TM5 | 7.98 | 0.08 | 9.10 | 0.16 |
| D275<sup>6.58</sup>N | TM6 | 7.73 | 0.04 | 8.79 | 0.06 |
| Q301<sup>7.39</sup>E | TM7 | 7.77 | 0.06 | 9.42 | 0.12 |
| Y51<sup>1.39</sup>A | TM1 | 7.31 | 0.08 | 8.61 | 0.08 |
| W100<sup>2.60</sup>Q | TM2 | no binding | - | 8.94 | 0.07 |
| H121<sup>3.29</sup>A | TM3 | 7.26 | 0.01 | 9.44 | 0.08 |
| F124<sup>3.32</sup>A | TM3 | no binding | - | 8.64 | 0.06 |
| V217<sup>5.43</sup>A | TM5 | 7.42 | 0.01 | 9.27 | 0.04 |
| Y268<sup>6.51</sup>A | TM6 | no binding | - | 9.67 | 0.02 |
| H298<sup>7.36</sup>A | TM7 | 7.35 | 0.09 | 9.63 | 0.13 |

Next, a second round of mutagenesis was performed in order to detail the binding of VUF15485 to hACKR3. To probe the two hydrophobic subpockets in the 7TM crevice of chemokine receptors (the so-called major and minor pockets (Arimont et al., 2017, Allen et al., 2007)), the hydrophobic and/or aromatic residues Y51^1.39^, W100^2.60^, H121^3.29^F124^5.42^, V217^5.43^, Y268^6.51^ and H298^7.36^ were mutated to smaller, non-aromatic residues (Table 2). Mutation of W100^2.60^Q, F124^5.42^A, and Y268^6.51^A fully abolished the binding of [^3^H]VUF15485, whereas no major changes in affinity were observed for the other mutants (Table 2). All tested mutants were able to bind CXCL12 with comparable affinities to the WT receptor, with the exception of the Y268^6.51^A mutant, which showed an increased affinity for CXCL12 (Table 2).

Subsequently, VUF15485 was docked into the cryo-EM structure of ACKR3 in complex with the partial agonist CCX662 (PDB entry 7SK9) (Yen et al., 2022). The binding poses were ranked and evaluated according to their binding scores and their adherence to the site-directed mutagenesis data. The proposed binding pose for VUF15485 is shown in Fig. 7 where the basic amine of the pyrrolidine ring of VUF15485 forms a salt-bridge to the acidic sidechain of D179^4.60^. The fluorophenyl moiety is placed within the deep hydrophobic subpocket of the major binding pocket, in between TM3 to TM7, facilitating a pi-stacking network between F129^3.37^, W265^6.48^, H269^6.52^ and potentially Y268^6.48^. The trimethoxy-phenyl group stacks with F124^3.32^ in the minor binding pocket located between TM1, 2, 3 and 7.

**Figure 7.**
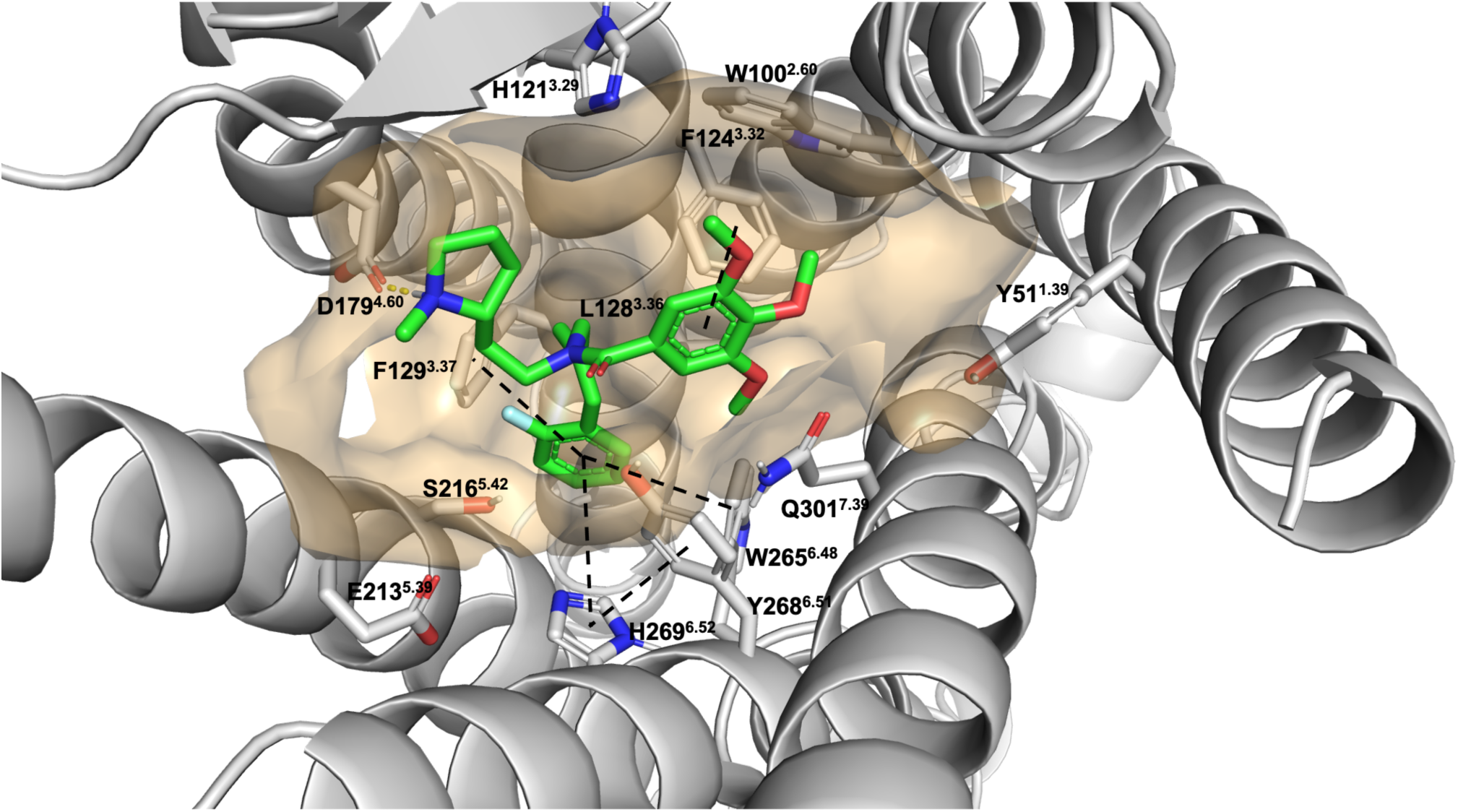
Docking pose of VUF15485 to hACKR3. VUF15485 was docked in the cryo-EM structure of hACKR3 (Yen et al., 2022) bound to the partial agonist CCX662 (PDB entry 7SK9). The side chains of selected residues are shown.

## Discussion

The purpose of this study was to synthesize and pharmacologically characterize the different enantiomers of a well-established small-molecule ACKR3 agonist, VUF11207 (Wijtmans et al., 2012a) in order to ultimately radiolabel the best enantiomer and characterize the first small-molecule agonist radioligand for an atypical chemokine receptor. We therefore synthesized both enantiomers, VUF13744 and VUF15485, and identified VUF15485 as the enantiomer with the highest affinity for hACKR3 (Fig.1). Moreover, in both a β−arrestin2 recruitment assay and a live cell microscopy setup with a new FLAsH-CFP-based conformational ACKR3 FRET sensor (Fig. 2), VUF15485 acts as an effective ACKR3 agonist, as reported previously for VUF11207 (Wijtmans et al., 2012a). Moreover, when tested against a panel of 20 human (classical and atypical) chemokine receptors, VUF15485 (1 µM) shows to be selective for ACKR3 and to not activate or antagonize other chemokine receptors (Fig. 3). Following these observations, it was decided to radiolabel the enantiopure VUF15485 by the incorporation of a CT_3_ group in order to generate and characterize a new research tool to study ACKR3.

The binding of [^3^H]VUF15485 to membranes of hACKR3-expressing HEK293T cells is saturable, occurs with high affinity and shows a clear pharmacological profile of ACKR3. A number of CXCR4 or CXCR3 ligands (in-house or established tool compounds) displace [^3^H]VUF15485 binding from hACKR3 only at high concentrations, whereas small-molecule ACKR3 ligands displace the binding with nanomolar affinities (Table 1). For example, in [^3^H]VUF15485 competition binding studies for the ACKR3 agonist VUF11403 a pIC_50_ value of 7.9 ± 0.1 was obtained, a similar value compared to what was published using [^125^I]CXCL12 on whole cells (p*K*_i_ = 7.7 ± 0.1) (Wijtmans et al., 2012a). Interestingly, [^3^H]VUF15485 binding to ACKR3 expressed in membranes of HEK293T cells is not displaced by the two endogenous ACKR3 ligands, CXCL11 and CXCL12. These data are in line with the observation that small-molecule agonists and the chemokines are targeting different ACKR3 conformations (Kleist et al., 2022). Moreover, previously similar observations have also been reported for radiolabeled small molecules binding to CXCR2 (de Kruijff et al., 2009) and CXCR3 (Scholten et al., 2015).

From kinetic association and dissociation experiments it can be determined that [^3^H]VUF15485 binding to hACKR3 occurs with very rapid kinetics leading e.g. to a very short residence time (RT) of < 2 min at 25°C. The short RT contrasts markedly with other published small-molecule radioligands targeting chemokine receptors, i.e. maraviroc, a CCR5 antagonist (RT > 136 hrs at 5°C) (Swinney et al., 2014), SCH527123, a CXCR2 antagonist (RT = 22 hrs) (Gonsiorek et al., 2007) and VUF11211, a CXCR3 antagonist (RT = 50 min) (Scholten et al., 2015). The rapid kinetics observed in the [^3^H]VUF15485 binding studies is in line with the rapid kinetics of VUF15485-induced conformational changes in ACKR3, as observed using a newly developed ACKR3-FlAsH-CFP conformational sensor. Previously, the insertion of FlAsH and CFP tags at appropriate intracellular locations in GPCR proteins has been used to register agonist-induced conformational changes of a number of GPCRs (Stumpf and Hoffmann, 2016). Following improved membrane localization of the ACKR3 sensor by the co-expression of a dominant negative K44A dynamin mutant, the ACKR3-FlAsH-CFP sensor responds rapidly to CXCL12 and VUF15485 and is the first successful report of a conformational sensor for a chemokine receptor. Next to the enantiopure, potent and selective ACKR3 agonist VUF15485 and the tritiated variant [^3^H]VUF15485, this new ACKR3 conformational sensor offers new opportunities for detailed pharmacological and biochemical studies of ACKR3.

In general, radioligand or fluorescence-based binding studies are key in drug discovery as they allow the accurate determination of equilibrium binding affinity constants and/or the kinetics of binding of GPCR ligands (Bayrak et al., 2022, Hopkins et al., 2022, Richard-Bildstein et al., 2020, Wijtmans et al., 2012a). Moreover, huge progress in the field of ACKR3 biochemistry has enabled NMR approaches with ^13^CH_3_-ε-methionine labelled ACKR3 and the elucidation of cryo-EM structures of antibody-stabilized ACKR3 with CXCL12, a CXCL12 analog and the ACKR3 agonist CCX662 (Kleist et al., 2022, Yen et al., 2022). These studies utilize different structural biology approaches and both indicate that the employed ACKR3 agonists (small-molecules CCX662 or CCX777, peptide LIH383) occupy distinct binding sites at ACKR3. Close comparison of the cryo-EM structures of ACKR3 with CXCL12 or CCX662 indicates that the small-molecule agonist occupies the orthosteric pocket within the transmembrane (TM) domains and mimics part of the chemokine N-terminus, but also occupies a distinct subpocket within the TM bundles (Yen et al., 2022). To actually provide detailed insights in the actual amino acids involved in the binding of available and new ACKR3 ligands, suitable labelled fluorescent and radioactive ligands, such as [^3^H]VUF15485, are a great asset as they allow site-directed mutagenesis studies to complement the available structural biology data. The current mutagenesis data (Table 2) shows abolished binding of VUF15485 at the W100^2.60^Q, F124^3.32^A, D179^4.60^N, and Y268^6.51^A mutants, highlighting the importance of these residues for ligand binding. Our computational modelling efforts suggests that the F124^3.32^A mutation disrupts the proposed pi-stacking between the aromatic residue and the trimethoxy-phenyl of VUF15485, while the D179^4.60^N mutation removes the negative charge, albeit retaining a hydrogen bond donor/acceptor character for this residue, supporting our observation of a direct ionic interaction between D179^4.60^ and the pyrrolidine group of VUF15485. W100^2.60^ forms a pi-stacking network with F124^3.32^ and nearby aromatic residues, stabilizing the contact of F124^3.32^ to VUF15485. The W100Q mutant, however, disrupts this network and introduces a bulkier and more hydrophilic residue likely promoting steric clashes with the trimethoxy-phenyl group and abolishing the binding of VUF15485.

In conclusion, VUF15485 has been pharmacologically characterized as the active enantiomer of the racemic ACKR3 agonist VUF11207. The compound proves to be a selective and potent ACKR3 agonist in a number of assays, including a newly developed ACKR3 conformational sensor. Following successful radiolabeling, [^3^H]VUF15485 was subsequently characterized in kinetic and equilibrium binding assays, revealing binding to hACKR3 with fast kinetics (i.e. a short residence time) and high affinity. Site-directed mutagenesis studies in combination with docking studies provide also a binding mode of the agonist in the ACKR3 binding site. [^3^H]VUF15485 is the first small-molecule radioligand for ACKR3 that can be used for detailed pharmacological investigations of ACKR3 and e.g. the evaluation of new molecules targeting ACKR3. Furthermore, the VUF15485 scaffold also offers interesting opportunities for the future development of other ACKR3 chemical biology tools, like e.g. fluorescent-labelled variants for optical imaging or ^18^F-labelled variants for Positron Emission Tomography.

## Acknowledgments

We thank Lonneke Rotteveel, Mark Stroet and Simon Mobach for technical assistance. This study was supported by the Luxembourg Institute of Health (LIH), Luxembourg National Research Fund (INTER/FNRS grants 20/15084569 and AFR HOPE-IOID, PRIDE-14254520 “I2TRON” and PRIDE-16749720 “NextImmune2”), F.R.S.-FNRS-Télévie (grants 7.8504.20, 7.4502.21 and 7.8508.22).

This research was funded by European Union’s Horizon2020 Marie Skłodowska-Curie Actions (MSCA) Program under Grant Agreement 641833 (ONCOgenic Receptor Network of Excellence and Training, ONCORNET) to AZ, IL, CP, CH, RL and IdE, MJS. RL, MS, CH, MW, HV, AC and MJS are part of the Marie Skłodowska-Curie Innovative Training Network ONCORNET2.0 “ONCOgenic Receptor Network of Excellence and Training” (MSCA-ITN-2020-ETN).

## References

Abagyan, R, Totrov, M, and Kuznetsov, D (1994) ICM—A New Method for Protein Modeling and Design: Applications to Docking and Structure Prediction from the Distorted Native Conformation. J Comput Chem, 15 (5), 488–506.

Adlere, I., Caspar, B., Arimont, M., Dekkers, S., Visser, K., Stuijt, J., de Graaf, C., Stocks, M., Kellam, B., Briddon, S., Wijtmans, M., de Esch, I., Hill, S., & Leurs, R. (2019). Modulators of CXCR4 and CXCR7/ACKR3 Function. Molecular Pharmacology, 96(6), 737–752.

Adlere I, Sun S, Zarca A, Roumen L, Gozelle M, Viciano CP, Caspar B, Arimont M, Bebelman JP, Briddon SJ, Hoffmann C, Hill SJ, Smit MJ, Vischer HF, Wijtmans M, de Graaf C, de Esch IJP and Leurs R (2019) Structure-based exploration and pharmacological evaluation of N-substituted piperidin-4-yl-methanamine CXCR4 chemokine receptor antagonists. Eur J Med Chem 162: 631–649.

Ainla A, Jansson ET, Stepanyants N, Orwar O and Jesorka A (2010) A microfluidic pipette for single-cell pharmacology. Anal Chem 82(11): 4529–4536.

Ainla A, Jeffries GD, Brune R, Orwar O and Jesorka A (2012) A multifunctional pipette. Lab Chip 12(7): 1255–1261.

Allen, S. J., Crown, S. E., & Handel, T. M. (2007). Chemokine: receptor structure, interactions, and antagonism. Annual Review of Immunology, 25, 787–820.

Arimont, M., Sun, S. L., Leurs, R., Smit, M., de Esch, I. J. P., & de Graaf, C. (2017). Structural Analysis of Chemokine Receptor-Ligand Interactions. Journal of Medicinal Chemistry, 60(12), 4735–4779.

Balabanian K, Lagane B, Infantino S, Chow KY, Harriague J, Moepps B, Arenzana-Seisdedos F, Thelen M and Bachelerie F (2005) The chemokine SDF-1/CXCL12 binds to and signals through the orphan receptor RDC1 in T lymphocytes. J Biol Chem 280(42): 35760–35766.

Ballesteros J. A. and Weinstein H. (1995) Integrated methods for the construction of three-dimensional models and computational probing of structure-function relations in G protein-coupled receptors. Methods in Neuroscience, 25, 366–428.

Bayrak, A., Mohr, F., Kolb, K., Szpakowska, M., Shevchenko, E., Dicenta, V., Rohlfing, A. K., Kudolo, M., Pantsar, T., Günther, M., Kaczor, A. A., Poso, A., Chevigné, A., Pillaiyar, T., Gawaz, M., & Laufer, S. A. (2022). Discovery and Development of First-in-Class ACKR3/CXCR7 Superagonists for Platelet Degranulation Modulation. Journal of Medicinal Chemistry, 65(19), 13365–13384.

Benredjem, B., Girard, M., Rhainds, D., St-Onge, G., & Heveker, N. (2017). Mutational Analysis of Atypical Chemokine Receptor 3 (ACKR3/CXCR7) Interaction with Its Chemokine Ligands CXCL11 and CXCL12. The Journal of Biological Chemistry, 292(1), 31–42.

Bobkov, V., Arimont, M., Zarca, A., De Groof, T. W. M., van der Woning, B., de Haard, H., & Smit, M. J. (2019). Antibodies Targeting Chemokine Receptors CXCR4 and ACKR3. Molecular Pharmacology, 96(6), 753–764.

Burns JM, Summers BC, Wang Y, Howard MC, Schall TJ and Miao Z, (2005) Methods and compositions for modulating angiogenesis, US 20050214287 A1.

Burns JM, Summers BC, Wang Y, Melikian A, Berahovich R, Miao Z, Penfold ME, Sunshine MJ, Littman DR, Kuo CJ, Wei K, McMaster BE, Wright K, Howard MC and Schall TJ (2006) A novel chemokine receptor for SDF-1 and I-TAC involved in cell survival, cell adhesion, and tumor development. J Exp Med 203(9): 2201–2213.

Canals M, Scholten DJ, de Munnik S, Han MK, Smit MJ and Leurs R (2012) Ubiquitination of CXCR7 controls receptor trafficking. PLoS One 7(3): e34192.

Cui, L. Y., Chu, S. F., & Chen, N. H. (2020). The role of chemokines and chemokine receptors in multiple sclerosis. International immunopharmacology, 83, 106314.

Dixon, A. S., Schwinn, M. K., Hall, M. P., Zimmerman, K., Otto, P., Lubben, T. H., Butler, B. L., Binkowski, B. F., Machleidt, T., Kirkland, T. A., Wood, M. G., Eggers, C. T., Encell, L. P., & Wood, K. V. (2016). NanoLuc Complementation Reporter Optimized for Accurate Measurement of Protein Interactions in Cells. ACS Chemical Biology, 11(2), 400–408.

Drouillard, D., Craig, B. T., & Dwinell, M. B. (2023). Physiology of chemokines in the cancer microenvironment. American Journal of Physiology. Cell Physiology, 324(1), C167–C182.

Flanagan C. A. (2016). GPCR-radioligand binding assays. Methods in cell biology, 132, 191–215.

Gonsiorek, W., Fan, X., Hesk, D., Fossetta, J., Qiu, H., Jakway, J., Billah, M., Dwyer, M., Chao, J., Deno, G., Taveras, A., Lundell, D. J., & Hipkin, R. W. (2007). Pharmacological characterization of Sch527123, a potent allosteric CXCR1/CXCR2 antagonist. The Journal of Pharmacology and Experimental Therapeutics, 322(2), 477–485.

Gustavsson M, Wang L, van Gils N, Stephens BS, Zhang P, Schall TJ, Yang S, Abagyan R, Chance MR, Kufareva I and Handel TM (2017) Structural basis of ligand interaction with atypical chemokine receptor 3. Nat Commun 8: 14135.

Hachet-Haas M, Balabanian K, Rohmer F, Pons F, Franchet C, Lecat S, Chow KY, Dagher R, Gizzi P, Didier B, Lagane B, Kellenberger E, Bonnet D, Baleux F, Haiech J, Parmentier M, Frossard N, Arenzana-Seisdedos F, Hibert M and Galzi JL (2008) Small neutralizing molecules to inhibit actions of the chemokine CXCL12. J Biol Chem 283(34): 23189–23199.

Hanes MS, Salanga CL, Chowdry AB, Comerford I, McColl SR, Kufareva I and Handel TM (2015) Dual targeting of the chemokine receptors CXCR4 and ACKR3 with novel engineered chemokines. J Biol Chem 290(37): 22385–22397.

Hatse S, Princen K, Liekens S, Vermeire K, De Clercq E and Schols D (2004) Fluorescent CXCL12AF647 as a novel probe for nonradioactive CXCL12/CXCR4 cellular interaction studies. Cytometry A 61(2): 178–188.

Hauser AS, Attwood MM, Rask-Andersen M, Schioth HB and Gloriam DE (2017) Trends in GPCR drug discovery: new agents, targets and indications. Nat Rev Drug Discov 16(12): 829–842.

Hoffmann C, Gaietta G, Bunemann M, Adams SR, Oberdorff-Maass S, Behr B, Vilardaga JP, Tsien RY, Ellisman MH and Lohse MJ (2005) A FlAsH-based FRET approach to determine G protein-coupled receptor activation in living cells. Nat Methods 2(3): 171–176.

Hoffmann C, Gaietta G, Zurn A, Adams SR, Terrillon S, Ellisman MH, Tsien RY and Lohse MJ (2010) Fluorescent labeling of tetracysteine-tagged proteins in intact cells. Nat Protoc 5(10): 1666–1677.

Hopkins, B. E., Masuho, I., Ren, D., Iyamu, I. D., Lv, W., Malik, N., Martemyanov, K. A., Schiltz, G. E., & Miller, R. J. (2022). Effects of Small Molecule Ligands on ACKR3 Receptors. Molecular Pharmacology, 102(3), 128–138.

Jost CA, Reither G, Hoffmann C and Schultz C (2008) Contribution of fluorophores to protein kinase C FRET probe performance. Chembiochem 9(9): 1379–1384.

Kalatskaya I, Berchiche YA, Gravel S, Limberg BJ, Rosenbaum JS and Heveker N (2009) AMD3100 is a CXCR7 ligand with allosteric agonist properties. Mol Pharmacol 75(5): 1240–1247.

Kleist, A. B., Jenjak, S., Sente, A., Laskowski, L. J., Szpakowska, M., Calkins, M. M., Anderson, E. I., McNally, L. M., Heukers, R., Bobkov, V., Peterson, F. C., Thomas, M. A., Chevigné, A., Smit, M. J., McCorvy, J. D., Babu, M. M., & Volkman, B. F. (2022). Conformational selection guides β-arrestin recruitment at a biased G protein-coupled receptor. Science, 377(6602), 222–228.

Kohler RE, Comerford I, Townley S, Haylock-Jacobs S, Clark-Lewis I and McColl SR (2008) Antagonism of the chemokine receptors CXCR3 and CXCR4 reduces the pathology of experimental autoimmune encephalomyelitis. Brain Pathol 18(4): 504–516.

de Kruijf, P., van Heteren, J., Lim, H. D., Conti, P. G., van der Lee, M. M., Bosch, L., Ho, K. K., Auld, D., Ohlmeyer, M., Smit, M. J., Wijkmans, J. C., Zaman, G. J., Smit, M. J., & Leurs, R. (2009). Nonpeptidergic allosteric antagonists differentially bind to the CXCR2 chemokine receptor. The Journal of Pharmacology and Experimental Therapeutics, 329(2), 783–790.

Lee, J., Cheng, X., Swails, J. M., Yeom, M. S., Eastman, P. K., Lemkul, J. A., Wei, S., Buckner, J., Jeong, J. C., Qi, Y., Jo, S., Pande, V. S., Case, D. A., Brooks, C. L., 3rd, MacKerell, A. D., Jr, Klauda, J. B., & Im, W. (2016). CHARMM-GUI Input Generator for NAMD, GROMACS, AMBER, OpenMM, and CHARMM/OpenMM Simulations Using the CHARMM36 Additive Force Field. Journal of Chemical Theory and Computation, 12(1), 405–413.

Libert, F., Parmentier, M., Lefort, A., Dumont, J. E., & Vassart, G. (1990). Complete nucleotide sequence of a putative G protein coupled receptor: RDC1. Nucleic acids research, 18(7), 1917.

Luker KE, Steele JM, Mihalko LA, Ray P and Luker GD (2010) Constitutive and chemokine-dependent internalization and recycling of CXCR7 in breast cancer cells to degrade chemokine ligands. Oncogene 29(32): 4599–4610.

Meiron M, Zohar Y, Anunu R, Wildbaum G and Karin N (2008) CXCL12 (SDF-1alpha) suppresses ongoing experimental autoimmune encephalomyelitis by selecting antigen-specific regulatory T cells. J Exp Med 205(11): 2643–2655.

Melikian, A, J. Burns, J., McMaster, B.E., Schall, T. and Wright, J.J. (2004) Inhibitors of human tumor-expressed CCXCKR2, WO20041058705 A2.

Meyrath, M., Szpakowska, M., Zeiner, J., Massotte, L., Merz, M. P., Benkel, T., Simon, K., Ohnmacht, J., Turner, J. D., Krüger, R., Seutin, V., Ollert, M., Kostenis, E., & Chevigné, A. (2020). The atypical chemokine receptor ACKR3/CXCR7 is a broad-spectrum scavenger for opioid peptides. Nature Communications, 11(1), 3033.

Mikolajczyk, T. P., Szczepaniak, P., Vidler, F., Maffia, P., Graham, G. J., & Guzik, T. J. (2021). Role of inflammatory chemokines in hypertension. Pharmacology & Therapeutics, 223, 107799.

Naumann U, Cameroni E, Pruenster M, Mahabaleshwar H, Raz E, Zerwes HG, Rot A and Thelen M (2010) CXCR7 functions as a scavenger for CXCL12 and CXCL11. PLoS One 5(2): e9175.

Neves, M. A., Totrov, M., & Abagyan, R. (2012). Docking and scoring with ICM: the benchmarking results and strategies for improvement. Journal of Computer-Aided Molecular Design, 26(6), 675–686.

Nibbs RJ and Graham GJ (2013) Immune regulation by atypical chemokine receptors. Nat Rev Immunol 13(11): 815–829.

Nijmeijer S, Engelhardt H, Schultes S, van de Stolpe AC, Lusink V, de Graaf C, Wijtmans M, Haaksma EE, de Esch IJ, Stachurski K, Vischer HF and Leurs R (2013) Design and pharmacological characterization of VUF14480, a covalent partial agonist that interacts with cysteine 98(3.36) of the human histamine H(4) receptor. Br J Pharmacol 170(1): 89–100.

Patel J, Channon KM and McNeill E (2013) The downstream regulation of chemokine receptor signalling: implications for atherosclerosis. Mediators Inflamm 2013: 459520.

Perpiñá-Viciano, C., Işbilir, A., Zarca, A., Caspar, B., Kilpatrick, L. E., Hill, S. J., Smit, M. J., Lohse, M. J., & Hoffmann, C. (2020). Kinetic Analysis of the Early Signaling Steps of the Human Chemokine Receptor CXCR4. Molecular Pharmacology, 98(2), 72–87.

Rajagopal S, Kim J, Ahn S, Craig S, Lam CM, Gerard NP, Gerard C and Lefkowitz RJ (2010) Beta-arrestin- but not G protein-mediated signaling by the “decoy” receptor CXCR7. Proc Natl Acad Sci U S A 107(2): 628–632.

Ray P, Mihalko LA, Coggins NL, Moudgil P, Ehrlich A, Luker KE and Luker GD (2012) Carboxy-terminus of CXCR7 regulates receptor localization and function. Int J Biochem Cell Biol 44(4): 669–678.

Richard-Bildstein, S., Aissaoui, H., Pothier, J., Schäfer, G., Gnerre, C., Lindenberg, E., Lehembre, F., Pouzol, L., & Guerry, P. (2020). Discovery of the Potent, Selective, Orally Available CXCR7 Antagonist ACT-1004-1239. Journal of Medicinal Chemistry, 63(24), 15864–15882.

Saaber F, Schutz D, Miess E, Abe P, Desikan S, Ashok Kumar P, Balk S, Huang K, Beaulieu JM, Schulz S and Stumm R (2019) ACKR3 Regulation of Neuronal Migration Requires ACKR3 Phosphorylation, but Not beta-Arrestin. Cell Rep 26(6): 1473–1488 e1479.

Sarma, P., Banerjee, R., & Shukla, A. K. (2023). Structural snapshot of a β-arrestin-biased receptor. Trends in Pharmacological Sciences, 44(1), 1–3.

Schols D, Este JA, Henson G and De Clercq E (1997) Bicyclams, a class of potent anti-HIV agents, are targeted at the HIV coreceptor fusin/CXCR-4. Antiviral Res 35(3): 147–156.

Scholten DJ, Canals M, Maussang D, Roumen L, Smit MJ, Wijtmans M, de Graaf C, Vischer HF and Leurs R (2012) Pharmacological modulation of chemokine receptor function. Br J Pharmacol 165(6): 1617–1643.

Scholten DJ, Roumen L, Wijtmans M, Verkade-Vreeker MC, Custers H, Lai M, de Hooge D, Canals M, de Esch IJ, Smit MJ, de Graaf C and Leurs R (2014) Identification of overlapping but differential binding sites for the high-affinity CXCR3 antagonists NBI-74330 and VUF11211. Mol Pharmacol 85(1): 116–126.

Scholten DJ, Wijtmans M, van Senten JR, Custers H, Stunnenberg A, de Esch IJ, Smit MJ and Leurs R (2015) Pharmacological characterization of [3H]VUF11211, a novel radiolabeled small-molecule inverse agonist for the chemokine receptor CXCR3. Mol Pharmacol 87(4): 639–648.

Shao Y, Anilkumar GN, Carroll CD, Dong G, Hall JW, 3rd, Hobbs DW, Jiang Y, Jenh CH, Kim SH, Kozlowski JA, McGuinness BF, Rosenblum SB, Schulman I, Shih NY, Shu Y, Wong MK, Yu W, Zawacki LG and Zeng Q (2011) II. SAR studies of pyridyl-piperazinyl-piperidine derivatives as CXCR3 chemokine antagonists. Bioorg Med Chem Lett 21(5): 1527–1531.

Shimizu S, Brown M, Sengupta R, Penfold ME and Meucci O (2011) CXCR7 protein expression in human adult brain and differentiated neurons. PLoS One 6(5): e20680.

Stumpf AD and Hoffmann C (2016) Optical probes based on G protein-coupled receptors - added work or added value? Br J Pharmacol 173(2): 255–266.

Sun X, Cheng G, Hao M, Zheng J, Zhou X, Zhang J, Taichman RS, Pienta KJ and Wang J (2010) CXCL12 / CXCR4 / CXCR7 chemokine axis and cancer progression. Cancer Metastasis Rev 29(4): 709–722.

Swinney, D. C., Beavis, P., Chuang, K. T., Zheng, Y., Lee, I., Gee, P., Deval, J., Rotstein, D. M., Dioszegi, M., Ravendran, P., Zhang, J., Sankuratri, S., Kondru, R., & Vauquelin, G. (2014). A study of the molecular mechanism of binding kinetics and long residence times of human CCR5 receptor small molecule allosteric ligands. British Journal of Pharmacology, 171(14), 3364–3375.

Szpakowska, M., Nevins, A. M., Meyrath, M., Rhainds, D., D’huys, T., Guité-Vinet, F., Dupuis, N., Gauthier, P. A., Counson, M., Kleist, A., St-Onge, G., Hanson, J., Schols, D., Volkman, B. F., Heveker, N., & Chevigné, A. (2018). Different contributions of chemokine N-terminal features attest to a different ligand binding mode and a bias towards activation of ACKR3/CXCR7 compared with CXCR4 and CXCR3. British Journal of Pharmacology, 175(9), 1419–1438.

Szpakowska, M., Meyrath, M., Reynders, N., Counson, M., Hanson, J., Steyaert, J., & Chevigné, A. (2018). Mutational analysis of the extracellular disulphide bridges of the atypical chemokine receptor ACKR3/CXCR7 uncovers multiple binding and activation modes for its chemokine and endogenous non-chemokine agonists. Biochemical Pharmacology, 153, 299–309.

Thoma G, Streiff MB, Kovarik J, Glickman F, Wagner T, Beerli C and Zerwes HG (2008) Orally bioavailable isothioureas block function of the chemokine receptor CXCR4 in vitro and in vivo. J Med Chem 51(24): 7915–7920.

Torphy, R. J., Yee, E. J., Schulick, R. D., & Zhu, Y. (2022). Atypical chemokine receptors: emerging therapeutic targets in cancer. Trends in Pharmacological Sciences, 43(12), 1085– 1097.

Ulvmar MH, Hub E and Rot A (2011) Atypical chemokine receptors. Exp Cell Res 317(5): 556–568.

Uto-Konomi A, McKibben B, Wirtz J, Sato Y, Takano A, Nanki T and Suzuki S (2013) CXCR7 agonists inhibit the function of CXCL12 by down-regulation of CXCR4. Biochem Biophys Res Commun 431(4): 772–776.

Wang C, Chen W and Shen J (2018) CXCR7 Targeting and Its Major Disease Relevance. Front Pharmacol 9: 641.

Wijtmans M, Maussang D, Sirci F, Scholten DJ, Canals M, Mujic-Delic A, Chong M, Chatalic KL, Custers H, Janssen E, de Graaf C, Smit MJ, de Esch IJ and Leurs R (2012a) Synthesis, modeling and functional activity of substituted styrene-amides as small-molecule CXCR7 agonists. Eur J Med Chem 51: 184–192.

Wijtmans M, Scholten DJ, Roumen L, Canals M, Custers H, Glas M, Vreeker MC, de Kanter FJ, de Graaf C, Smit MJ, de Esch IJ and Leurs R (2012b) Chemical subtleties in small-molecule modulation of peptide receptor function: the case of CXCR3 biaryl-type ligands. J Med Chem 55(23): 10572–10583.

Wijtmans M, Verzijl D, Bergmans S, Lai M, Bosch L, Smit MJ, de Esch IJ and Leurs R (2011) CXCR3 antagonists: quaternary ammonium salts equipped with biphenyl- and polycycloaliphatic-anchors. Bioorg Med Chem 19(11): 3384–3393.

Wurth R, Bajetto A, Harrison JK, Barbieri F and Florio T (2014) CXCL12 modulation of CXCR4 and CXCR7 activity in human glioblastoma stem-like cells and regulation of the tumor microenvironment. Front Cell Neurosci 8: 144.

Yen, Y. C., Schafer, C. T., Gustavsson, M., Eberle, S. A., Dominik, P. K., Deneka, D., Zhang, P., Schall, T. J., Kossiakoff, A. A., Tesmer, J. J. G., & Handel, T. M. (2022). Structures of atypical chemokine receptor 3 reveal the basis for its promiscuity and signaling bias. Science Advances, 8(28), eabn8063.

Verweij, E. W. E., Al Araaj, B., Prabhata, W. R., Prihandoko, R., Nijmeijer, S., Tobin, A. B., Leurs, R., & Vischer, H. F. (2020). Differential Role of Serines and Threonines in Intracellular Loop 3 and C-Terminal Tail of the Histamine H_4_ Receptor in β-Arrestin and G Protein-Coupled Receptor Kinase Interaction, Internalization, and Signaling. ACS Pharmacol & Translational Sci, 3(2), 321–333.

Zarca, A., Perez, C., van den Bor, J., Bebelman, J. P., Heuninck, J., de Jonker, R. J. F., Durroux, T., Vischer, H. F., Siderius, M., & Smit, M. J. (2021). Differential Involvement of ACKR3 C-Tail in β-Arrestin Recruitment, Trafficking and Internalization. Cells, 10(3), 618.

