## Supplementary information for "Pharmacological characterization and radiolabeling of VUF15485, a high-affinity small-molecule agonist for the atypical chemokine receptor ACKR3"

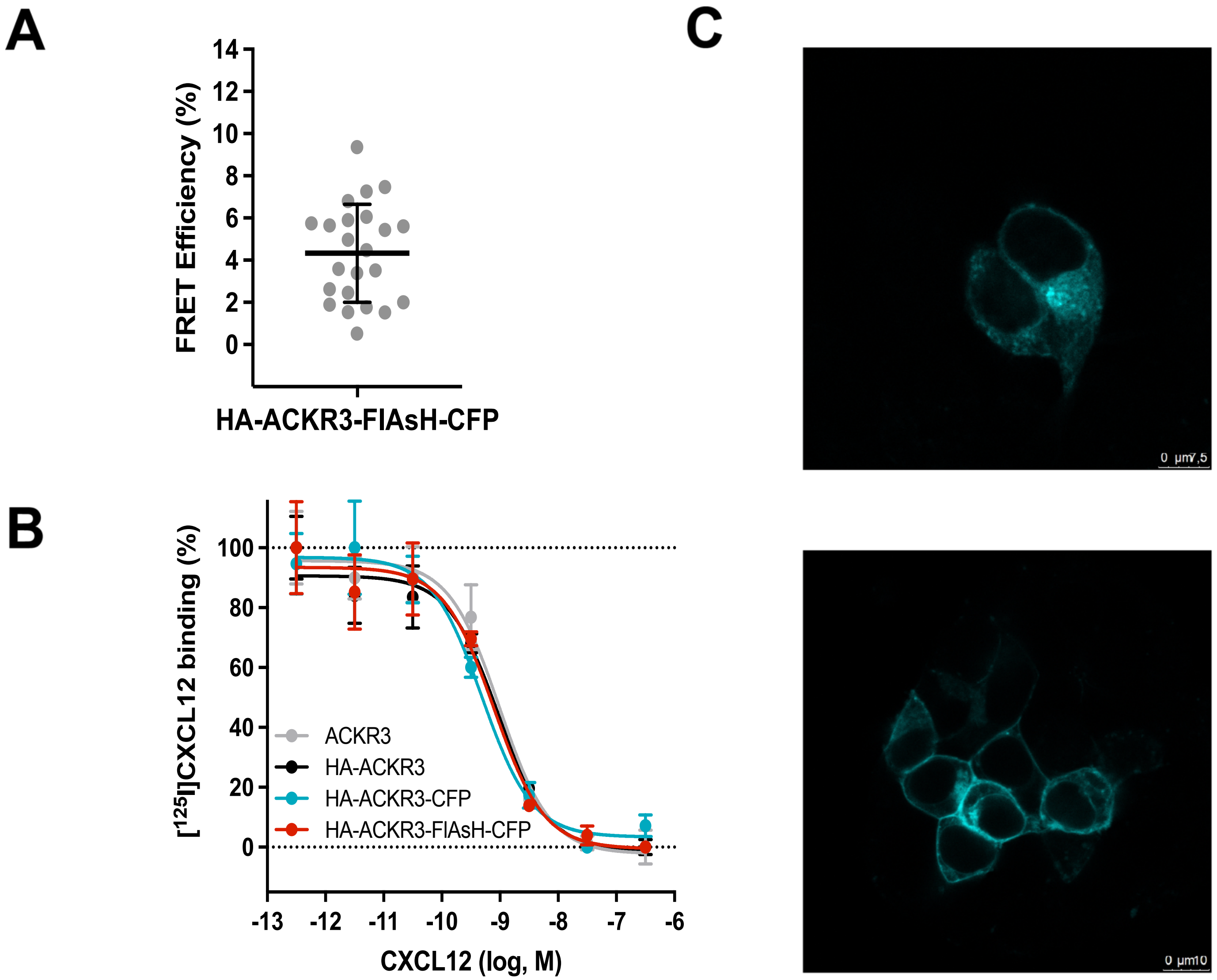


**Figure S1**. Characterization of a FRET-based hACKR3 conformational sensor. A) FRET efficiency of the HA-tagged hACKR3-FlAsH-CFP sensor was determined by BAL treatment to be 4.33 ± 2.33 %. Data shows mean ± SD of 23 cells measured in four independent experiments. No intermolecular FRET was detected (data not shown). B) CXCL12 binding to HA-tagged hACKR3, hACKR3-CFP and hACKR3-FlAsH-CFP. Competition binding was performed using [$^{125}$I]CXCL12 and increasing doses of CXCL12 as competitor. Representative curves with triplicates are plotted with SD. C) Confocal image of HEK293 cells transiently expressing the ACKR3-FlAsH-CFP construct alone (upper panel) or with K44A dynamin (lower panel). CFP emission is shown upon excitation at 442 nm. The scale bar represents 10 μm.

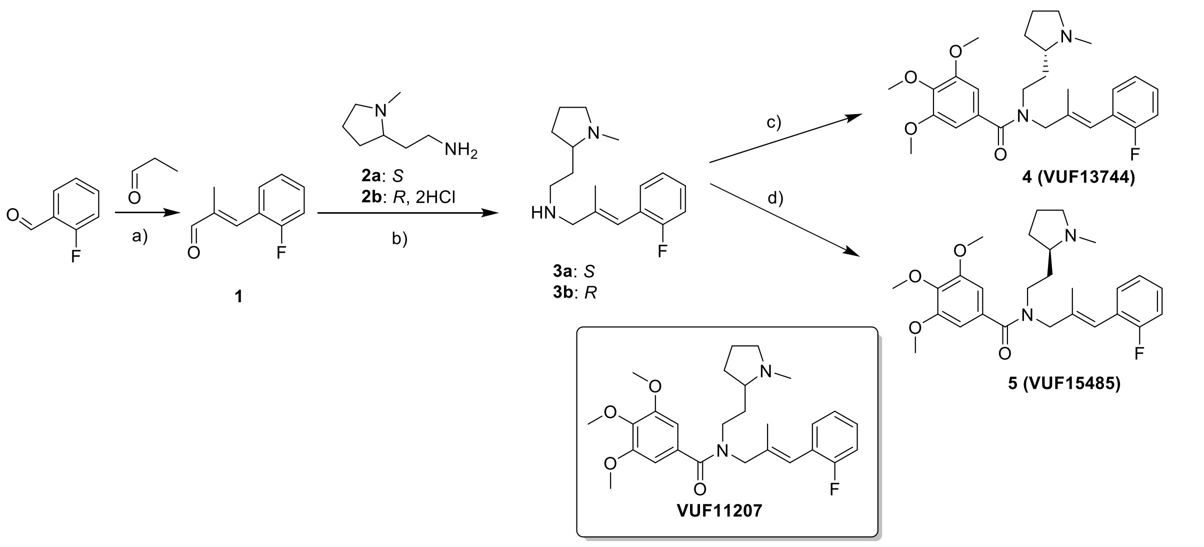


**Scheme S1. Synthesis of VUF11207 stereoisomers**. Reagents and conditions: a) KOH, $H_2O$, EtOH, 24 h, 0 °C → rt, 74 %; b) (i) $Na_2SO_4$, DCM, 24 h, rt, TEA for **2b**, (ii) $NaBH_4$, MeOH, 0.5 h, 0 °C, **3a**: 74%; **3b**: 62 %; ee > 95%; c) 3,4,5-trimethoxybenzoyl chloride, TEA, DCM, 16 h, rt, 29%; ee > 95%; d) 3,4,5-trimethoxybenzoic acid, HOBt hydrate, EDCI, DIPEA, THF, 40 h, rt, 62%; ee > 95%.

Enantiopure isomers of VUF11207 were prepared by starting with the respective enantiopure 2-(1-methylpyrrolidin-2-yl)ethan-1-amine **2** according to our previously disclosed route (Scheme S1) (Wijtmans et al, 2012). An indirect reductive amination of aldehyde **1** (prepared from 2-fluorobenzaldehyde in 74 % yield) with enantiopure amine **2a** or **2b** yielded enantiopure key intermediates **3a** and **3b** in good yields (74% and 62%, respectively). Imine formation was monitored with $^{1}H$ NMR spectroscopy for a maximum conversion before the reduction with $NaBH_4$. Chiral liquid chromatography (LC) analysis of amines **3a,b** gave completely resolved enantiomer peaks and, given the availability of racemic **3** in our labs, allowed determination of enantiomeric excess (ee) for

**3a,b** to be > 95 %. Acylation of amine **3a** with 3,4,5-trimethoxybenzoyl chloride, prepared from $SOCl_2$ and the corresponding benzoic acid, resulted in the *S*-enantiomer of VUF11207 (i.e. VUF13744, **4**) in 29% yield. For the *R*-enantiomer (VUF15485, **5**) we resorted to a higher-yielding procedure using EDCI/HOBt hydrate mediated amide coupling yielding **5** in good yield (62%). Despite extensive efforts, chiral LC analysis on the final products **4** and **5** did not allow completely resolved peaks, although the chiral LC analysis qualitatively suggests high ee and it is assumed that the ee of precursors **3a,b** would not erode during the final amidation step.

**Synthesis of $^3$H-labelled analogs of VUF11207**

The scaffold of VUF11207 provides multiple methyl groups that could be considered for late-stage $CT_3$ incorporation. Introduction of a $CT_3$ group on the amine was discarded due to low availability of suitable enantiopure precursors. Rather, substitution of one of the methoxy moieties is preferred, especially since required chemicals are commercially available. With the purpose of evaluating the preferred position for labelling, we prepared phenol analogues of VUF11207 (compounds **8** and **9**) with the hydroxy group at the *para*- and *meta*-position, respectively. Racemic precursors **8** and **9** were prepared via amide couplings as described for **5** in moderate yields (31% and 37%, respectively) and subjected to trial labelings with [$^3$H]methylnosylate (Scheme S2). As Figure S2 shows, *meta*-compound **9** showed higher reactivity towards labeling within the time course analysed. We attribute this to steric and electronic factors that hinder the labelling of precursor **8**.

Based on these data, the 3-hydroxy unit (as in **9**) was chosen for the production of [$^3$H]VUF15485 using the *R*-enantiomer of **9**, compound **6** (Scheme S3), resulting in *(3-[$^3$H]methoxy)*VUF15485. Precursor **6** was prepared similarly to racemate **9** with an amide coupling using enantiopure amine **3b**. Radiolabeled *(3-[$^3$H]methoxy)*VUF15485 was obtained from methylation of phenol **6** with [$^3$H]methylnosylate in ACN in the presence of aqueous NaOH solution (Scheme S3).

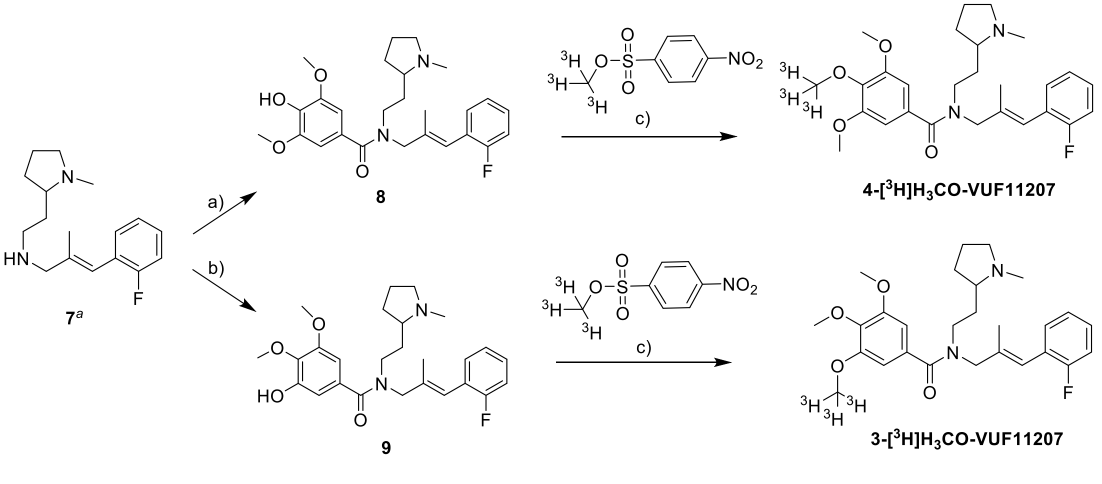


**Scheme S2. Synthesis and trial radiolabeling of racemic precursors.** Reagents and conditions: a) 4-hydroxy-3,5-dimethoxybenzoic acid, HOBt hydrate, EDCI, DIPEA, THF, 24 h, rt, 31%; b) 3-hydroxy-4,5-dimethoxybenzoic acid, HOBt hydrate, EDCI, DIPEA, THF, 36 h, rt, 37%; c) NaOH, MeCN, 25 °C, timepoints at 20, 40, 60, 140 or 220 min (see Fig. S2). [a]Compound **7** was prepared and characterized as reported (Wijtmans et al, 2012).

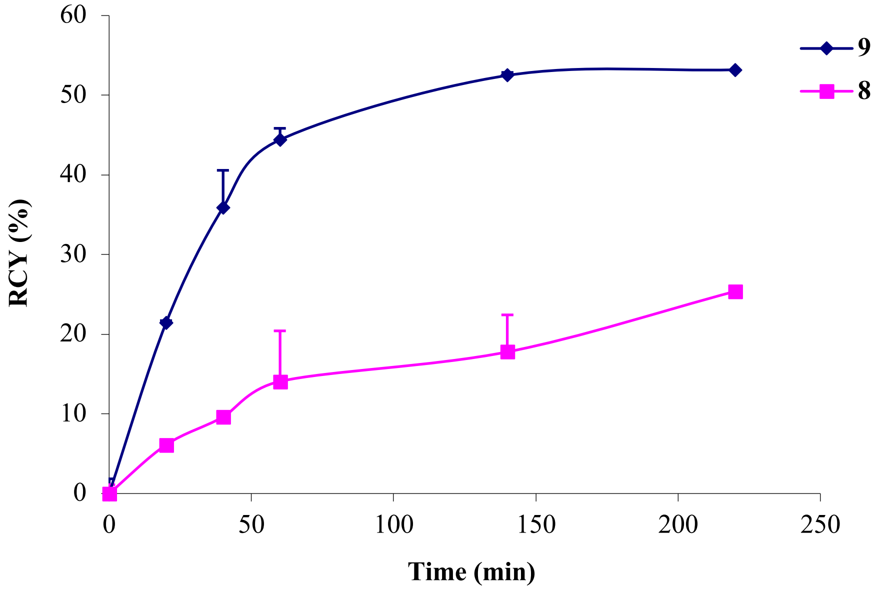


**Figure S2.** Conversion of precursors **9** and **8** to (*3-[$^3$H]methoxy*)VUF11207 and (*4-[$^3$H]methoxy*)VUF11207, respectively, as measured at different reaction time points at 25 °C. The radiochemical yield (RCY in %) was determined based on the HPLC with radioactivity detection. Two independent experiments were carried out and the average values with SD are shown.

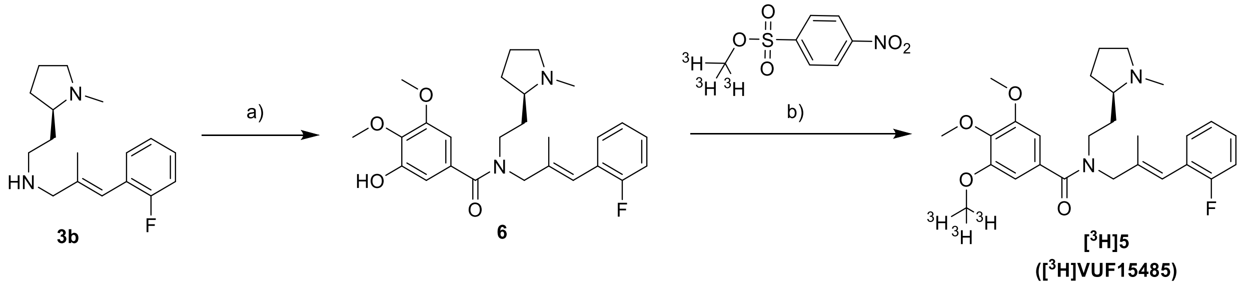


**Scheme S3. Radiosynthesis of [$^3$H]VUF15485**. Reagents and conditions: a) 3-hydroxy-4,5-dimethoxybenzoic acid, HOBt hydrate, EDCI, DIPEA, THF, 24 h, rt, 33%; b) NaOH, MeCN, 60° C, 40 min, RCY = 38%.

The scaffold of **5** provides multiple methyl groups that could be considered for late-stage $CT_3$ incorporation. However, installing a $CT_3$ group on the amine was discarded due to low availability of suitable enantiopure precursors. Rather, introducing a $CT_3$ unit on one of the two possible phenols was deemed a better approach. With the purpose of evaluating the preferred position for the radiolabeling of the phenols, we prepared phenol analogues of VUF11207 (compounds **8** and **9**) with the hydroxy group at *para*- and *meta*- positions, respectively. Racemic precursors **8** and **9** were prepared via amide couplings as described for **5** in moderate yields (31% and 37%, respectively) and subjected to trial labelling with [$^3$H]methylnosylate (Scheme S2). As Figure S2 shows, *meta*-compound **9** showed higher reactivity towards labeling within the time course analysed. We attribute this to steric and electronic factors that are more severe with precursor **8**. Thus, the 3-hydroxy unit (as in **9**) was chosen for the production of [$^3$H]VUF15485 using the *R*-enantiomer of **9**, i.e. **6** (Scheme S3). Enantiomerically pure precursor **6** was prepared similarly to racemate **9** with an amide coupling using enantiopure amine **3b**. Radiolabeled [$^3$H]VUF15485 was obtained from methylation of phenol **6** with [$^3$H]methylnosylate in ACN in the presence of aqueous NaOH solution (Scheme S3). To increase the conversion rate, heating to 60˚C was applied. This yielded a total amount of 38 MBq (38% RCY, radiochemical purity of 98%). Determination of the ee of *(3-[$^3$H]methoxy)*VUF15485 was hampered by its low amount of mass and hence UV signal and by inefficient baseline separation on chiral LC analysis on precursor **6**. To remedy this, we mimicked the methylation of **6** with a non-radiolabelled methylation reagent (e.g. MeI//$K_2CO_3$/$CH_3COCH_3$/60 min at reflux → 14 h at rt) and measured the ee of the resulting alkylated product, qualitatively providing an ee of > 95% with the correct mass. This indicates that the compound **6** was sufficiently enantiopure and by extension this is assumed to hold true for *(3-[$^3$H]methoxy)*VUF15485 as well, given that during radiolabeling the ee is unlikely to erode.

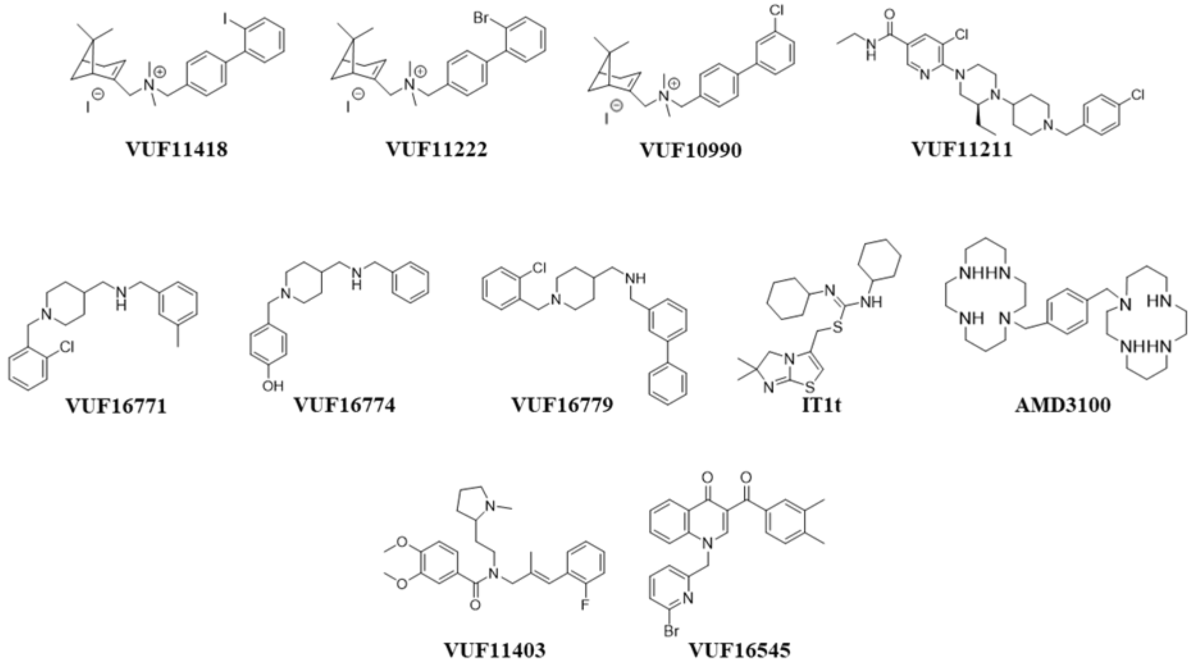


**Figure S3.** Structures of selected unlabeled CXCR3, CXCR4 and ACKR3 chemokine receptor small-molecule ligands.

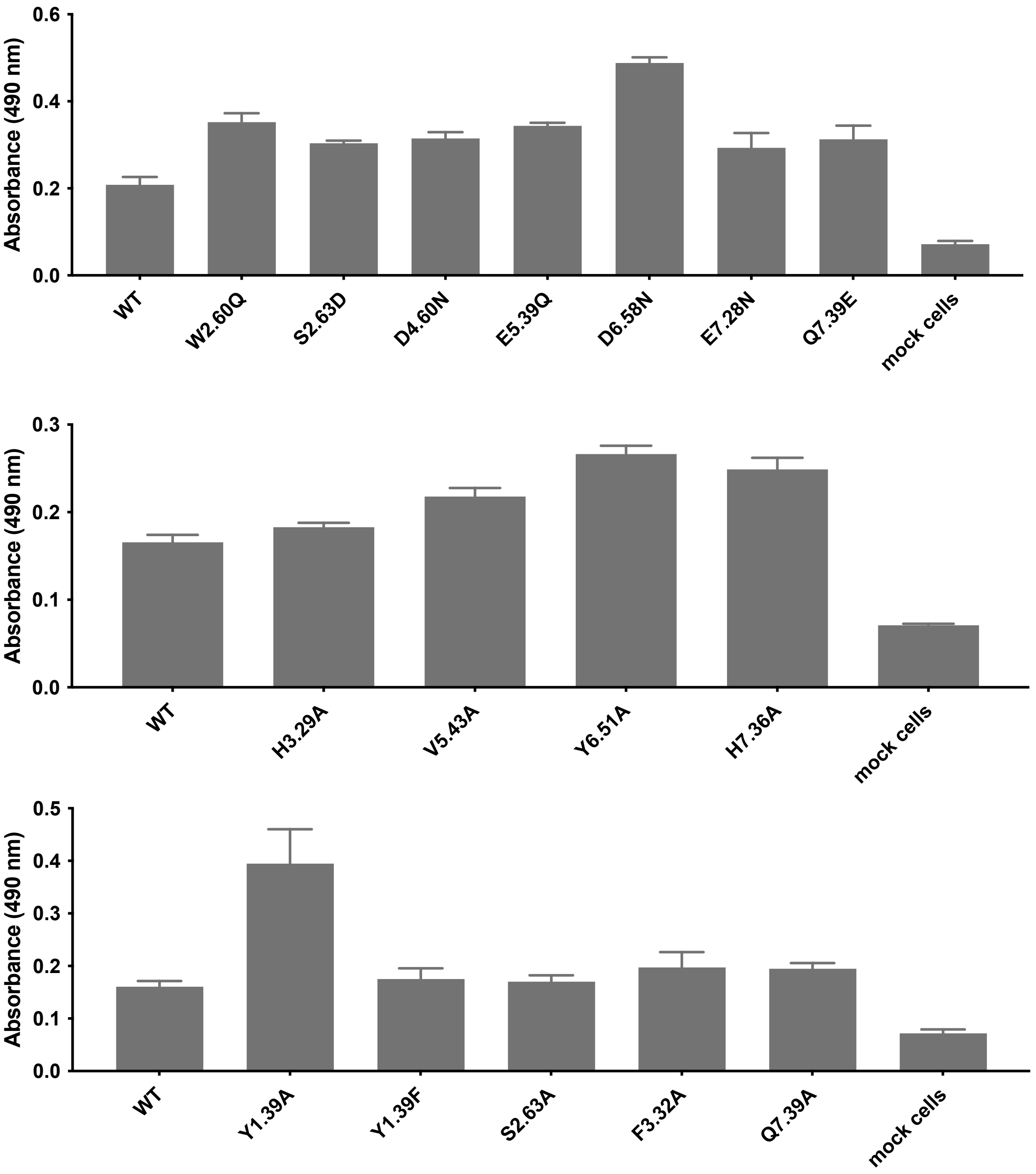


**Figure S4 Expression of HA-tagged wild type (WT) and mutant hACKR3 proteins upon transient expression in HEK293-T cells.** Receptor proteins were detected on cell membranes by anti-HA ELISA as described previously (Verweij et al., 2020).

**Synthetic chemistry procedures**

Dry solvents (THF, DCM) were obtained from a PureSolv solvent purification system (Inert®). All other commercial reagents and solvents were used without further purification. Enantiopure amines **2a** and **2b** (diHCl salt) were obtained from Astatech and Activate Scientific, respectively. All reactions were carried out under an inert $N_2$ atmosphere unless otherwise stated. TLC analyses were performed with Merck F254 Alumina Silica Plates using UV visualization or staining. Column purifications were carried out automatically using Isolera One Biotage® equipment. $^{1}H$ and $^{13}C$ spectra were recorded on a Bruker spectrometer with operating frequency 400 MHz, 500 MHz, 600 MHz and 101 MHz, 126 MHz, 151 MHz respectively. NMR spectra were calibrated according to internal references for non-deuterated solvents: $CHCl_3$ ($\delta_H$ = 7.26 ppm), $CDCl_3$ ($\delta_C$ = 77.16 ppm), DMSO ($\delta_H$ = 2.50), DMSO-$d_6$ ($\delta_C$ = 39.52 ppm) and $H_2O$ ($\delta_H$ = 4.79). The following abbreviations are used to denote multiplicities: s = singlet, d = doublet, t = triplet, q = quartet, m = multiplet, br = broad signal, app = apparent. Systematic names for molecules according to IUPAC rules were generated using ChemDraw Pro 16.0. All HRMS spectra were recorded on Bruker microTOF-Q MS using ESI in positive ion mode. The purity of compounds was determined using a Shimadzu HPLC/MS workstation with a LC-20AD pump system, SPD-M20A diode array detection and a LCMS-2010 EV Liquid Chromatograph Mass Spectrometer (for LR-MS) and applying either a basic or acidic mode. Compound purities were calculated as the percentage peak area of the analyzed compound by UV detection at, unless stated otherwise, 254 nm. The column used is an Xbridge C18 5 µm column (50 mm×4.6 mm). *Basic mode:* Solvent B (MeCN/10% buffer), Solvent A ($H_2O$/10% buffer). The buffer is a 0.4% (w/v) $NH_4HCO_3$ solution in $H_2O$, adjusted to pH 8.0 with $NH_4OH$. The analysis was conducted using a flow rate of 1.0 mL/ min with a total run time of 8 min or 12 min depending on the lipophilicity of the analyte. *Acidic mode*: Solvent B (MeCN/0.1% formic acid) and solvent A ($H_2O$/0.1% formic acid), flow rate of 1.0 mL/min with a run time of 8 min. Gradient settings: For 8 min run (basic and acidic system): start 5% B, linear gradient to 90% B in 4.5 min, then isocratic for 1.5 min at 90% B, then linear gradient to 5% B in 0.5 min, then isocratic for 1.5 min at 5% B. For 12 min run (basic system): start 5% B, linear gradient to 90% B in 4.5 min, then 5.5 min at 90% B, then linear gradient to 5% B in 0.5 min, then isocratic for 1.5 min at 5% B. Normal phase chiral LC analyses were performed with system 1, 2 or 3. *System 1*: Shimadzu LC-20AD liquid chromatograph pump system, equipped with a Phenomenex® Lux™ Cellulose-1 column (5 µm, 250x4.6 mm), connected to a Shimadzu SPD-M20A diode array detector (254 nm) and using an isocratic approach (heptane / MeOH / DEA 90 : 10: 0.1 ) and flow rate of 0.5 mL/min with a run time of 15 min. *System 2:* Shimadzu VP Series HPLC using a diode array detector with a Chiralcel OD-H (150x4.6 mm) column and heptane / 2-propanol 97 : 3 as eluent at 1.0 ml/min. Chiral supercritical fluid chromatography (SFC) analysis (*System 3*) were performed on Shimadzu NexeraUC LC-30AD liquid chromatograph pump system, equipped with Phenomenex Lux Cellulose-1 column (5 µm, 250x4.6 mm), connected to Shimadzu SPD-M20A diode array detector, Shimadzu NexeraUC SFC-30A back pressure regulator and MS detection using a Shimadzu LC-MS-2020 mass spectrometer. Conditions applied were an isocratic system ($CO_2$ / MeOH / DEA 95 : 5: 0.5 ) and flow rate of 1.0 mL/min with a run time of 60 min.

***Synthesis***

**(*E*)-3-(2-Fluorophenyl)-2-methylacrylaldehyde (1)**

The synthetic procedure reported by us (Wijtmans et al., 2012a) was followed, using 2-fluorobenzaldehyde (6.95 g, 56.00 mmol), 1-propanal (3.39 g, 58.40 mmol), KOH (0.31 g, 5.60 mmol), 95% EtOH (45 mL), $H_2O$ (0.4 mL) and a reaction time of 24 h. Compound **1** was obtained as a yellow oil (6.82 g, 74 %) with LC purity of 92%. The compound was used without further purification. The spectral data correspond to the data reported (Wijtmans et al., 2012a). $^1$H NMR

($CDCl_3$, 500 MHz) δ 9.63 (s, 1H), 7.58-7.49 (m, 1H), 7.48-7.34 (m, 2H), 7.26-7.11 (m, 2H), 2.03 (s, 3H).

**(*S,E*)-3-(2-fluorophenyl)-2-methyl-N-(2-(1-methylpyrrolidin-2-yl)ethyl)prop-2-en-1-amine (3a)**

Amine **2a** (0.26 g, 2.00 mmol), aldehyde **1** (0.33 g, 2.00 mmol) and $Na_2SO_4$ (1.70 g, 12.00 mmol) were mixed with DCM (10 mL). The mixture was stirred at room temperature for 24 h and imine formation was followed by $^1$H NMR spectroscopy of concentrated samples. The reaction mixture was filtered and the collected solid was washed with DCM. The combined filtrates were concentrated *in vacuo*. MeOH (10 mL) was added to the residue and the mixture was cooled to 0 °C. $NaBH_4$ (0.100 g, 2.64 mmol) was slowly added to reduce the imine. The mixture was stirred at 0 °C for 30 min. The reaction mixture was quenched with acetone (1 mL) and $H_2O$ (10 mL), stirred for 10 min at room temperature and concentrated *in vacuo*. $H_2O$ (25 mL) was added to the residue and extraction with DCM (2x25 mL) was performed, ensuring that the pH of the aq. layer is >10 by addition of aq. 10% $K_2CO_3$ solution. The combined organic layers were washed with brine (1x), dried over $Na_2SO_4$, filtered and concentrated *in vacuo*. The crude product was purified by column chromatography (Biotage) with eluent system EtOAc:MeOH:TEA (80 : 20 : 2). Compound **3a** was obtained as a light-yellow oil (0.40 g, 74%). $^1$H NMR (400 MHz, $CDCl_3$) δ 7.28-6.98 (m, 5H), 6.41 (s, 1H), 3.35 (s, 2H), 3.11-2.99 (m, 1H), 2.76-2.58 (m, 2H), 2.32 (s, 3H), 2.17-2.06 (m, 2H), 1.96-1.61 (m, 8H), 1.53-1.43 (m, 2H). ESI-MS *m/z:* 277.00 $[M + H]^+$. LC purity: 96% (230 nm). ee > 95% (*System 2*).

**(*R,E*)-3-(2-fluorophenyl)-2-methyl-N-(2-(1-methylpyrrolidin-2-yl)ethyl)prop-2-en-1-amine (3b)**

Amine dihydrochloride **2b** (0.35 g, 1.73 mmol), aldehyde **1** (0.28 g, 1.73 mmol), TEA (0.35 g, 3.46 mmol) and $Na_2SO_4$ (1.42 g, 10.00 mmol) were mixed with DCM (10 mL). The mixture was stirred at room temperature for 24 hours and imine formation was followed by $^1$H NMR spectroscopy of concentrated samples. The reaction mixture was filtered and the collected solid was washed with DCM. The combined filtrates were concentrated *in vacuo*. MeOH (10 mL) was added to the residue and the mixture was cooled to 0 °C. $NaBH_4$ (0.07 g, 1.875 mmol) was slowly added to reduce the imine. The mixture was stirred at 0 °C for 30 min. The reaction mixture was quenched with acetone (1 mL) and $H_2O$ (10 mL), stirred for 10 min at room temperature and concentrated *in vacuo*. $H_2O$ (25 mL) was added to the residue and extraction with DCM (2x25 mL) was performed, ensuring that the pH of the aq. layer is >10 by addition of 10% aqueous $K_2CO_3$ solution. The combined organic layers were washed with brine (1x), dried over $Na_2SO_4$, filtered and concentrated *in vacuo*. The crude product was purified by column chromatography (Biotage) with eluent system EtOAc:MeOH/10% TEA (10% to 25%). Compound **3b** was obtained as a light-yellow oil (0.30 g, 62%). $^1$H NMR (400 MHz, $CDCl_3$) δ 7.30-6.94 (m, 4H), 6.40 (s, 1H), 3.34 (s, 2H), 3.11-2.96 (m, 1H), 2.78-2.55 (m, 2H), 2.32 (s, 3H), 2.15-2.03 (m, 2H), 2.03-1.60 (m, 8H), 1.58-1.38 (m, 2H). HRMS-ESI *m/z* $[M + H]^+$ calc. for $C_{17}H_{26}FN_2^+$ 277.2075; found 277.2064. LC purity: 97% (254 nm). ee > 95% (*System 1*).

**(*S,E*)-N-(3-(2-fluorophenyl)-2-methylallyl)-3,4,5-trimethoxy-N-(2-(1-methylpyrrolidin-2-yl)ethyl)benzamide (4, VUF13744)**

3,4,5-Trimethoxybenzoic acid (1.07 g, 5.04 mmol) was mixed with $SOCl_2$ (16.3 g, 0.137 mol). The mixture was heated at reflux for 2 h. $SOCl_2$ was removed *in vacuo*. The solid was dissolved in DCE (20 mL) and the mixture was concentrated *in vacuo* (2x). This afforded the acid chloride, which was immediately used in the crude form. Amine **3a** (0.24 g, 0.868 mmol) was mixed with TEA (0.200 g, 2.276 mmol) in anhydrous DCM (8 mL). To this, a solution of the crude acid chloride (0.360 g) in DCM (2 mL) was added dropwise at rt. The mixture was stirred for 16 h at room temperature and then made basic with saturated aqeous $Na_2CO_3$ (10 mL). The organic layer was dried over $Na_2SO_4$, filtered and concentrated *in vacuo*. The crude product was purified by two sequential column chromatography purifications (Biotage) with eluent system EtOAc:MeOH:TEA (80 : 20 : 2) and EtOAc:heptane:TEA (90 : 10 : 1), respectively. This gave **4** as a colorless oil (0.12 g, 29%). $^1$H NMR

(400 MHz, DMSO-$d_6$, 373 K) δ 7.31-7.10 (m, 4H), 6.65 (s, 2H), 6.36 (s, 1H), 4.09 (s, 2H), 3.77 (s, 6H), 3.70 (s, 3H), 3.34 (t, 2H, $J$ = 8.0 Hz), 3.00-2.85 (m, 1H, overlaps with HDO peak), 2.15 (s, 3H), 2.10-1.95 (m, 2H), 1.94-1.70 (m, 2H), 1.69 (s, 3H), 1.62-1.55 (m, 3H), 1.40-1.20 (m, 1H). $^{13}$C NMR (101 MHz, DMSO-$d_6$, 373 K) δ 170.8, 161.0, 153.5, 139.9, 137.6, 132.7, 130.9, 129.2, 124.5, 124.5, 119.3, 115.7, 105.6, 64.1, 60.6, 57.0, 56.9, 41.1, 40.4, 31.5, 30.3, 22.3, 16.2. ESI-MS *m/z:* 471.10 [M + H]$^+$. HRMS-ESI *m/z* [M + H]$^+$ calc. for $C_{27}H_{36}FN_2O_4^+$ 471.2654; found 471.2635. LC purity: 98% (254 nm). ee > 95% (*System 3*).

**General procedure for EDCI-mediated amide coupling**

EDCI (1.2 eq) and DIPEA (2.5 eq) were added to a mixture of amine **3a**, **3b** or **7**, HOBt hydrate (1.2 eq) and the corresponding benzoic acid (1.0 eq) in THF. The mixture was stirred at room temperature for the time indicated after which the reaction mixture was quenched with aq. 10% $K_2CO_3$ to reach a pH value of ca. 11. Extraction was performed with $H_2O$ (20 mL) and EtOAc (2x20 mL). The combined organic layers were washed with brine (1x), dried over $MgSO_4$ and concentrated *in vacuo*. The crude product was purified by column chromatography (Biotage) with eluent system EtOAc:MeOH/50% TEA (0 to 10%).

**(*R,E*)-N-(3-(2-fluorophenyl)-2-methylallyl)-3,4,5-trimethoxy-N-(2-(1-methylpyrrolidin-2-yl)ethyl)benzamide (5, VUF15485)**

The general procedure EDCI-mediated amide coupling was followed using EDCI (0.16 g, 0.84 mmol), DIPEA (0.23 g, 1.76 mmol), amine **3b** (0.19 g, 0.70 mmol), HOBt (0.13 g, 0.84 mmol), 3,4,5-trimethoxybenzoic acid (0.15 g, 0.70 mmol), THF (20 mL) and a reaction time of 40 h. Compound **5** was obtained as a colorless oil (0.20 g, 62%). $^1$H NMR (400 MHz, DMSO-$d_6$, 373 K) δ 7.38-7.08 (m, 4H), 6.69 (s, 2H), 6.40 (s, 1H), 4.13 (s, 2H), 3.80 (s, 6H), 3.74 (s, 3H), 3.37 (t, $J$ = 7.8 Hz, 2H), 2.27-2.07 (m, 5H), 1.97-1.76 (m, 2H), 1.72 (s, 3H), 1.67-1.55 (m, 3H), 1.44-1.29 (m, 1H). $^{13}$C NMR (101

MHz, $CDCl_3$, 298 K) δ 171.8, 153.3, 136.3, 130.6, 128.7, 123.8, 115.7, 115.5, 103.9, 61.0, 57.2, 56.3, 43.0, 40.5, 40.3, 30.6, 29.8, 22.0, 16.2. HRMS-ESI *m/z* $[M + H]^+$ calc. for $C_{27}H_{36}FN_2O_4^+$ 471.2654; found 471.2635. LC purity: 98% (254 nm). ee > 95% (*System 3*).

**Synthesis of [$^3$H]VUF15485 ((*3-[$^3$H]methoxy*) VUF15485)**

***Precursors***

***Rac*-(*E*)-N-(3-(2-fluorophenyl)-2-methylallyl)-4-hydroxy-3,5-dimethoxy-N-(2-(1-methylpyrrolidin-2-yl)ethyl)benzamide (8)**

The general procedure EDCI-mediated amide coupling was followed using EDCI (0.12 g, 0.60 mmol), DIPEA (0.16 g, 1.25 mmol), amine **7** (0.14 g, 0.50 mmol), HOBt hydrate (0.09 g, 0.60 mmol), 4-hydroxy-3,5-dimethoxybenzoic acid (0.10 g, 0.50 mmol), THF (15 mL) and a reaction time of 24 h. Compound **8** was obtained as a light yellow oil (0.07 g, 31%). $^1$H NMR (400 MHz, DMSO-$d_6$, 373 K) δ 8.17 (s, 1H), 7.36-7.26 (m, 2H), 7.23-7.11 (m, 2H), 6.69 (s, 2H), 6.41 (s, 1H), 4.14 (s, 2H), 3.79 (s, 6H), 3.37 (t, *J* = 7.9 Hz, 2H), 2.93-2.87 (m, 1H), 2.20 (s, 3H), 2.11-2.01 (m, 2H), 1.95-1.75 (m, 2H), 1.71 (s, 3H), 1.66-1.52 (m, 3H), 1.41-1.29 (m, 1H). HRMS-ESI *m/z* $[M + H]^+$ calc. for $C_{26}H_{34}FN_2O_4^+$ 457.2497; found 457.2504. LC purity: 98% (254 nm).

***Rac*-(*E*)-N-(3-(2-fluorophenyl)-2-methylallyl)-3-hydroxy-4,5-dimethoxy-N-(2-(1-methylpyrrolidin-2-yl)ethyl)benzamide (9)**

The general procedure EDCI-mediated amide coupling was followed using EDCI (0.12 g, 0.60 mmol), DIPEA (0.16 g, 1.25 mmol), amine **7** (0.14 g, 0.50 mmol), HOBt hydrate (0.09 g, 0.60 mmol), 3-hydroxy-4,5-dimethoxybenzoic acid (0.10 g, 0.50 mmol), THF (15 mL) and a reaction time of 36 h. Compound **9** was obtained as a light yellow oil (0.08 g, 37%). $^1$H NMR (400 MHz, DMSO-$d_6$, 373 K) δ 8.95 (s, 1H), 7.37-7.25 (m, 2H), 7.22-7.11 (m, 2H), 6.55-6.47 (m, 2H), 6.38 (s, 1H), 4.13 (s, 2H), 3.78 (s, 3H), 3.74 (s, 3H), 3.34 (t, *J* = 7.9 Hz, 2H), 2.94-2.87 (m, 1H), 2.18 (s, 3H), 2.13-1.99 (m, 2H),

1.93-1.74 (m, 2H), 1.71 (s, 3H), 1.65-1.50 (m, 3H), 1.38-1.25 (m, 1H). HRMS-ESI *m/z* $[M + H]^+$ calc. for $C_{26}H_{34}FN_2O_4^+$ 457.2497; found 457.2511. LC purity: 97% (254 nm).

**(*R,E*)-N-(3-(2-fluorophenyl)-2-methylallyl)-3-hydroxy-4,5-dimethoxy-N-(2-(1-methylpyrrolidin-2-yl)ethyl)benzamide (6)**

The general procedure EDCI-mediated amide coupling was followed using EDCI (0.11 g, 0.59 mmol), DIPEA (0.16 g, 1.23 mmol), amine **3b** (0.14 g, 0.49 mmol), HOBt hydrate (0.09 g, 0.60 mmol), 3-hydroxy-4,5-dimethoxybenzoic acid (0.10 g, 0.49 mmol), THF (20 mL) and a reaction time of 24 h. Compound **6** was obtained as a light yellow oil (0.07 g, 33%). $^1$H NMR (400 MHz, DMSO-$d_6$, 373 K) δ 8.95 (s, 1H), 7.37-7.25 (m, 2H), 7.22-7.11 (m, 2H), 6.55-6.47 (m, 2H), 6.38 (s, 1H), 4.13 (s, 2H), 3.78 (s, 3H), 3.74 (s, 3H), 3.34 (t, *J* = 7.9 Hz, 2H), 2.94-2.87 (m, 1H), 2.18 (s, 3H), 2.13-1.99 (m, 2H), 1.93-1.74 (m, 2H), 1.71 (s, 3H), 1.65-1.50 (m, 3H), 1.38-1.25 (m, 1H). HRMS-ESI *m/z* $[M + H]^+$ calc. for $C_{26}H_{34}FN_2O_4^+$ 457.2497; found 457.2511. LC purity: 96% (254 nm). Chiral analysis on **6** gave inefficient baseline separation, but chiral LC analysis of the product obtained from methylation of **6** with MeI (1.0 eq MeI//$K_2CO_3$/$CH_3COCH_3$/60 min at reflux for 14 hours at room temperature) qualitatively showed an ee > 95 % (*System 3*), underscoring that also **6** was enantiopure.

***Radiolabeling general***

A stock solution of 1.0 GBq [$^3$H]methylnosylate with molar activity 2.98 TBq/mmol in ACN (1 mL) was aliquoted per 0.100 mL and used further for the radiolabelling. The test reactions and the final product were analysed with HPLC equipped with UV/Vis (UV2075 Plus Jasc, 254 nm) and Beta-RAM radiation detectors (LabLogic, Model 4). HPLC analysis was done using Gracesmart RP C18 5µm (250x4.6 mm) column with eluent ACN/$H_2O$/TFA 35:65:0.1 and flow rate 1 mL/min. HPLC purification was done using Luna C18 10 µm (250x10 mm) column and isocratic elution with (ACN/$H_2O$/TFA 35:65:0.1) and a flow rate of 4 mL/min. The radioactivity counts per second were measured at least three times of the sample of 3 μL dissolved in 20 mL of the scintillation liquid (Optiphase Hisafe 3).

***Trial radiolabelling of racemic precursors***

A stock solution was made of 7.5 mg/mL of non-labelled methylnosylate. The resulting stock solution (10 μL) was diluted with 980 μL of ACN and 10 μL of the [$^3$H]methylnosylate stock solution (1.0 GBq/mL). The resulting solution contained 10 MBq/mL of activity and equal molarity of methylnosylate (labelled and non-labelled). Precursor **8** or **9** (0.5 mg, 1.1 μmol) was dissolved in ACN (200 μL). To the resulting solution aqueous 5M NaOH solution (5 μL) was added, and subsequently 100 μL of the radioactive and non-radioactive methylnosylate stock solution was added. The HPLC measurements were taken by pipetting 3 μL of the reaction mixture and quenching in the HPLC eluent (ACN/$H_2O$/TFA:35:65:0.1).

**[$^3$H](*E*)-*N*-(3-(2-fluorophenyl)-2-methylallyl)-3,-[$^3$H]methoxy-4,5-dimethoxy-*N*-(2-(1-methylpyrrolidin-2-yl)ethyl)benzamide ([$^3$H]5, (*3-[$^3$H]methoxy*)VUF15485)**

[$^3$H]methylnosylate (100 MBq, 100 μL of stock solution in ACN) and 0.5 mg of the precursor **6** were dissolved in 200 μL ACN. Next, 5 μL 5M aq. NaOH was added. The reaction was allowed to proceed for 40 min at 60˚C. The mixture was quenched with 0.6 mL of HPLC eluent and purified by HPLC using a Luna C18 column. The product was eluted with 35/65 ACN/$H_2O$+0.1%TFA at 4 mL/min and detection at 254 nm. Collected fractions were diluted with sterile $H_2O$ (50 mL) and passed over a preconditioned (pre-eluted with 25 mL $H_2O$; 10 mL EtOH; 25 mL $H_2O$) solid phase extraction cartridge (Waters tC18 Sep-Pak), which was subsequently washed with 20 mL of $H_2O$ for injection. The product was then fractionally eluted two times with 0.250 mL and 0.414 mL of 96% EtOH to yield the stock solutions of the title compound with 33 MBq/mL and 72 MBq/mL concentrations, yielding total of 38 MBq (38% radiochemical yield). Chiral LC analysis of (*3-[$^3$H]methoxy*)VUF15485 was not performed, but the product obtained from methylation of precursor **6** with MeI (1.0 eq MeI//$K_2CO_3$/$CH_3COCH_3$/60 min at reflux and then for 14 hours at room temperature) showed an ee > 95 % (*System 3*), underscoring that precursor **6** was as expected enantiopure and by extension it is assumed, given the similarity of the methylation steps, that also (*3-[$^3$H]methoxy*)VUF15485 was sufficiently enantiopure.

### Spectra and chromatograms

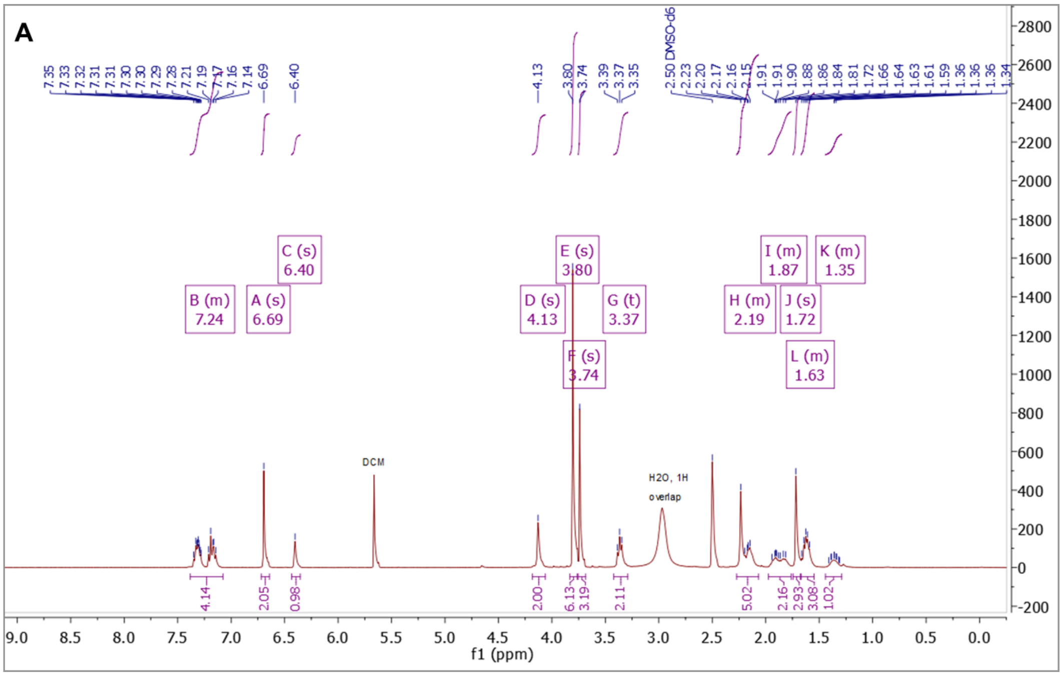


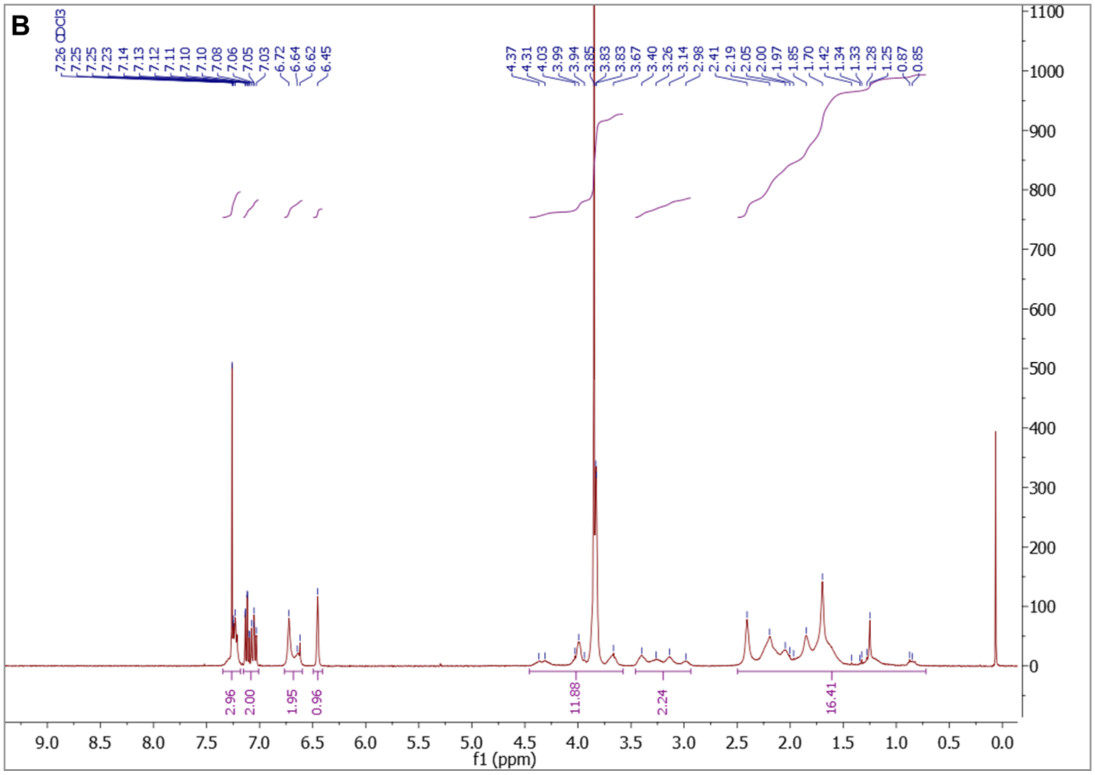

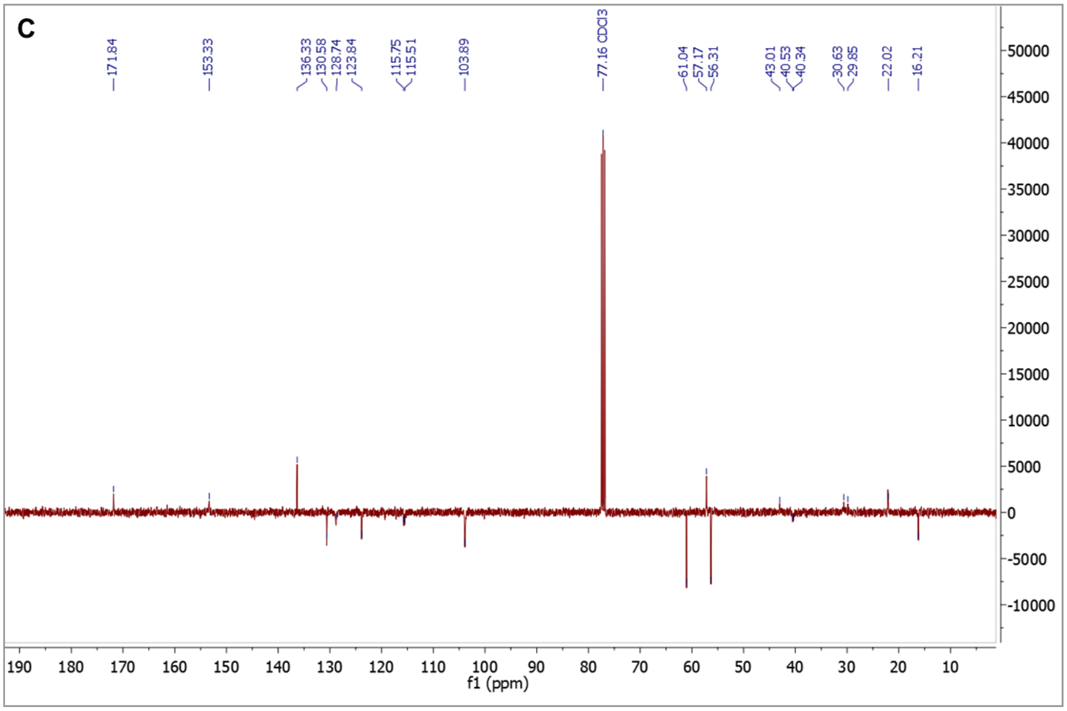


**Figure S5**. [1]H NMR (A: in DMSO-$d_6$ at 373 K at 400 MHz; B: in $CDCl_3$ at room temperature at 400 MHz) and [13]C NMR (C, in $CDCl_3$ at room temperature at 101 MHz) spectra of compound **5** (VUF15485). A trace of DCM is visible in the batch shown in Fig. A but this DCM was not present in the batch used for pharmacology (shown in Fig. B).

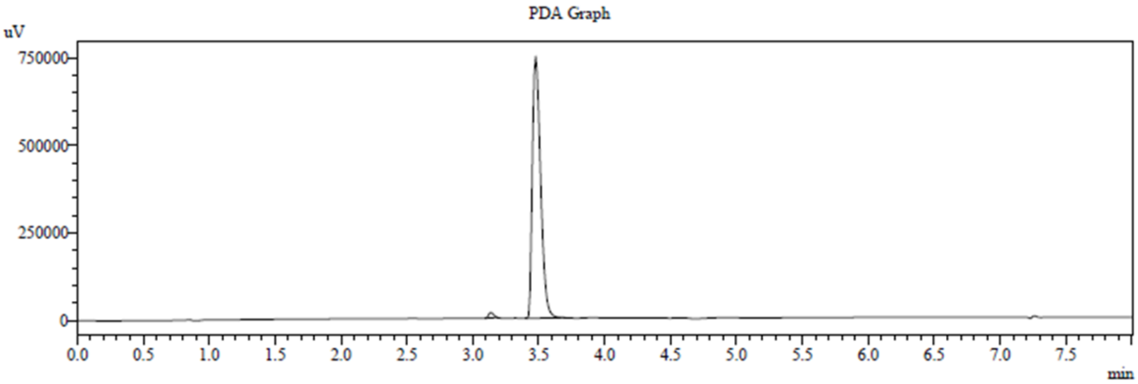


PDA Ch1 254nm 4nm

| Peak# | Name | Ret. Time | Area | Area % |
|---|---|---|---|---|
| 1 | | 3.132 | 42480 | 1.302 |
| 2 | | 3.472 | 3220020 | 98.698 |

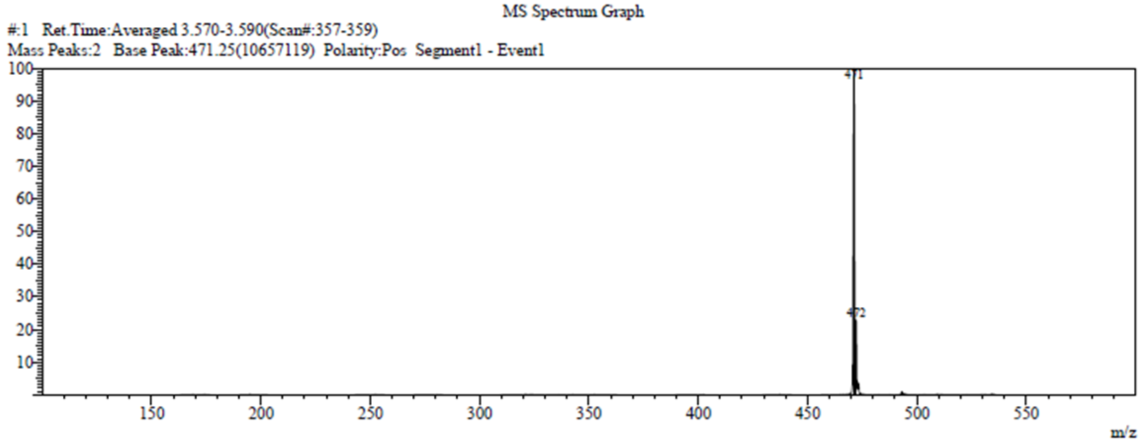


MS Spectrum Table

**Figure S6**. HPLC/MS chromatogram of compound **5** (VUF15485).

A

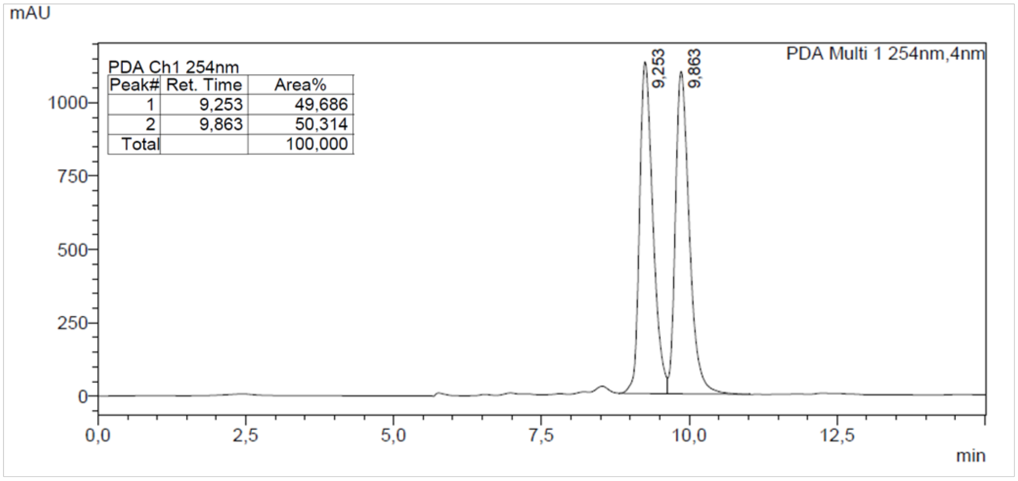


B

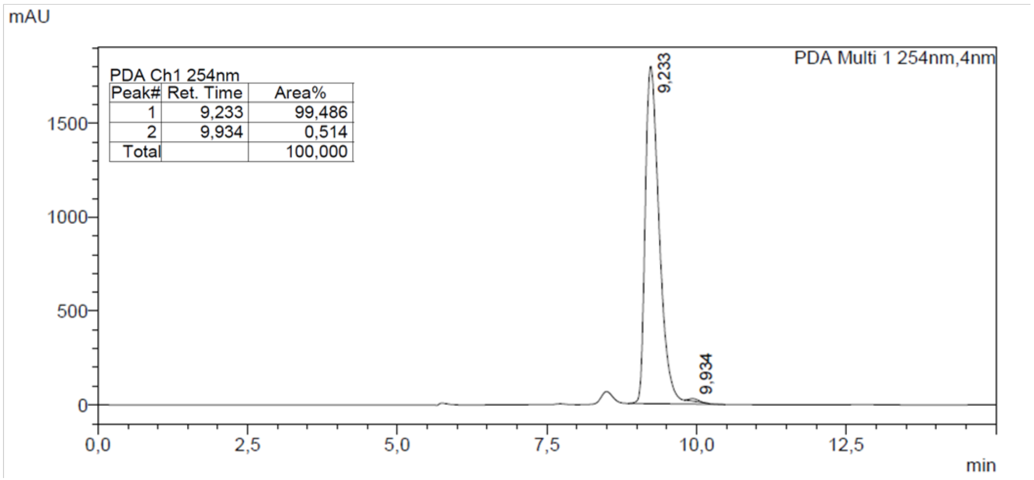


**Figure S7**. Chiral separation of racemic precursor **3** (A) and enantiomeric excess determination of *R*-enantiomer **3b** (B), measured with chiral normal phase HPLC *(System 1*) and detection at 254 nm.

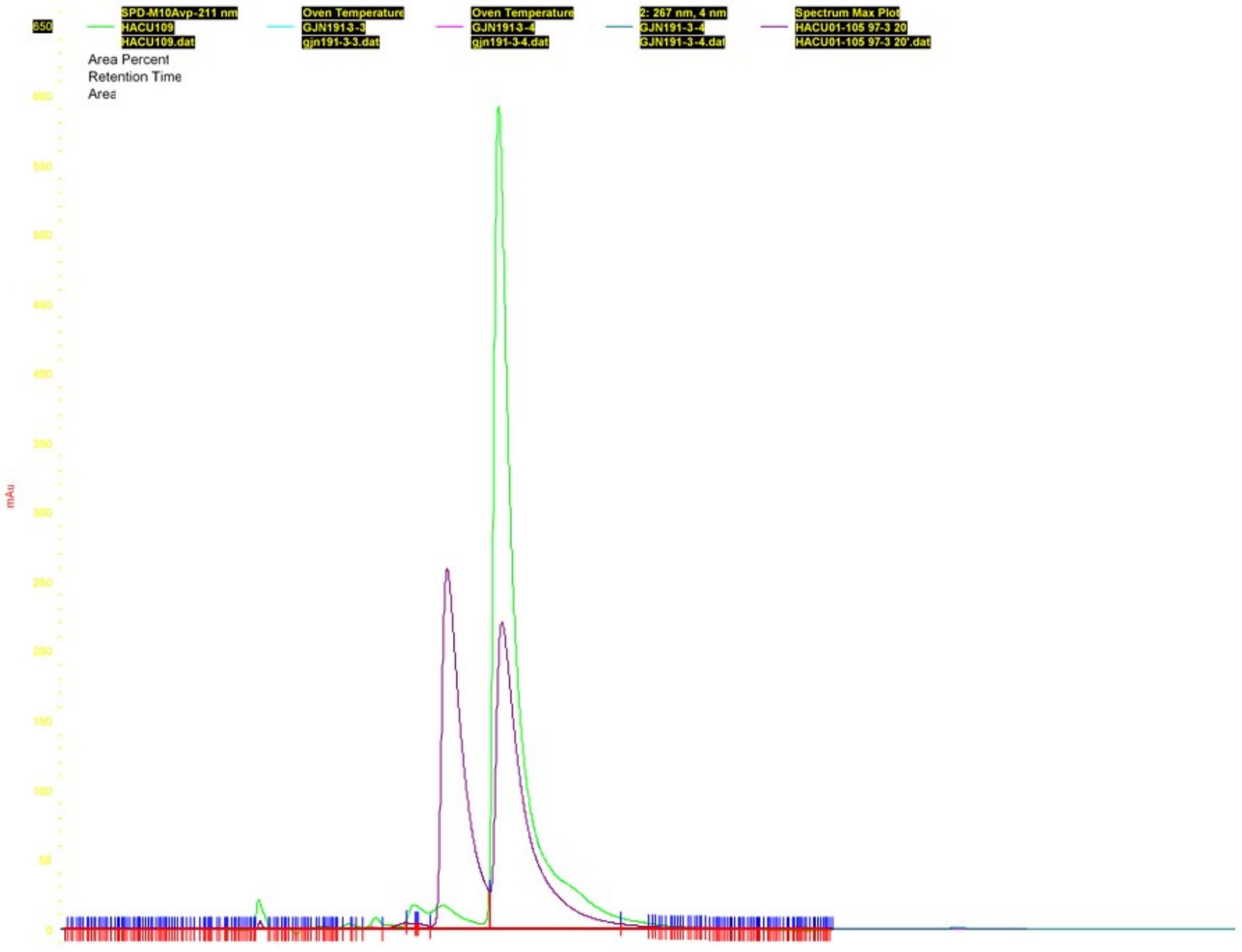


**Figure S8**. Chiral separation of racemic precursor **3** (purple) and enantiomeric excess determination of *S*-enantiomer **3a** (green), measured with chiral normal phase HPLC (*System 2*) and detection at 211 nm.

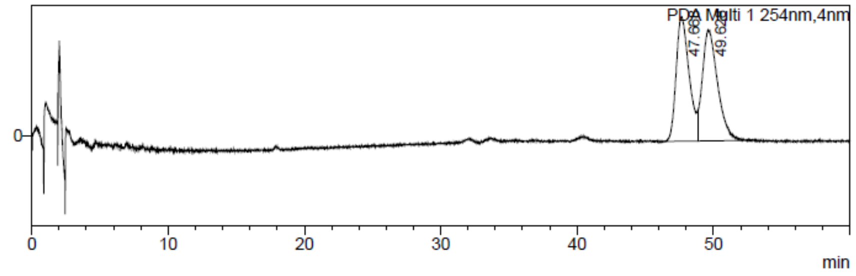


Peak Table

PDA Ch1 254nm

| Peak# | Ret. Time | Area% |
|---|---|---|
| 1 | 47.668 | 49.097 |
| 2 | 49.628 | 50.903 |
| Total | | 100.000 |

MS Chromatogram

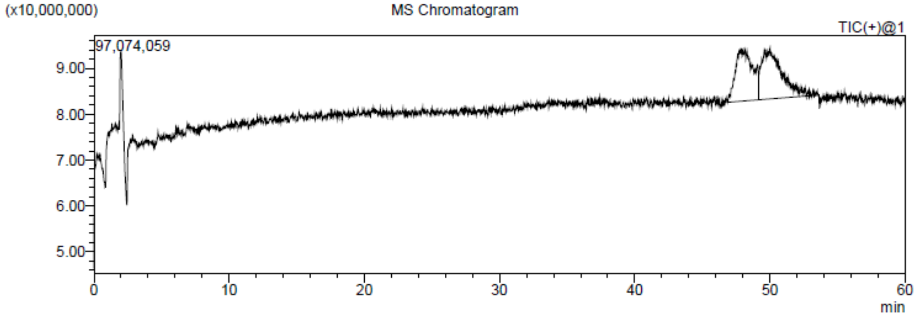


MASS Peak Table TIC

| Peak# | Ret. Time | m/z |
|---|---|---|
| 1 | 47.953 | TIC |
| 2 | 49.996 | TIC |
| Total | | |

MS Spectrum

Line#:1 R.Time:----(Scan#:----)
MassPeaks:6
Spectrum Mode:Averaged 47.817-48.233(2870-2895) Base Peak:471(9025342)
BG Mode:Averaged 20.033-22.033(1203-1323) Segment 1 - Event 1

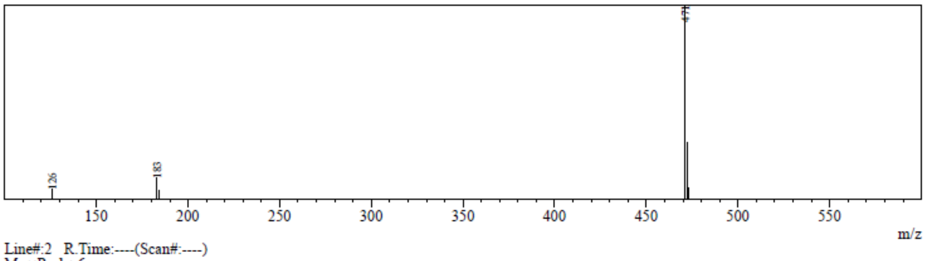


Line#:2 R.Time:----(Scan#:----)
MassPeaks:6
Spectrum Mode:Averaged 49.767-50.167(2987-3011) Base Peak:471(9276155)
BG Mode:Averaged 20.033-21.967(1203-1319) Segment 1 - Event 1

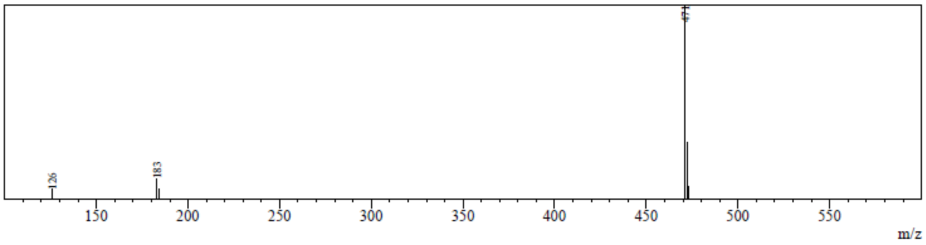


**Figure S9**. Chiral separation of racemic VUF11207 (i.e. **4** and **5**) measured with SFC/MS (*System 3*) and detection at 254 nm.

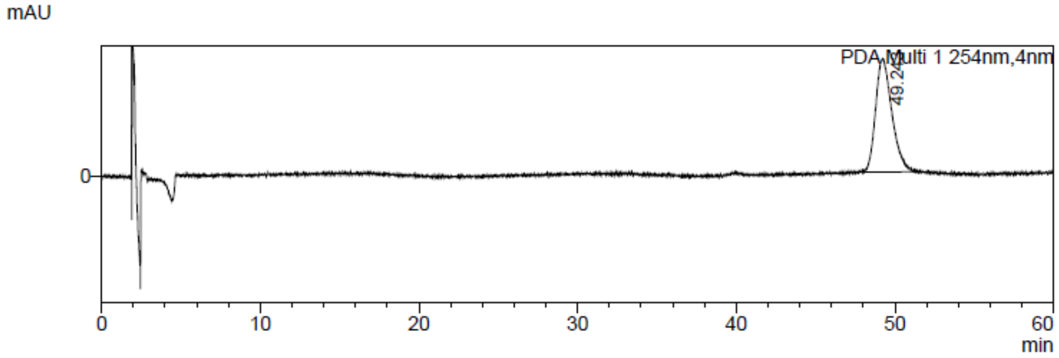


Peak Table

PDA Ch1 254nm

| Peak# | Ret. Time | Area% |
|---|---|---|
| 1 | 49.243 | 100.000 |
| Total | | 100.000 |

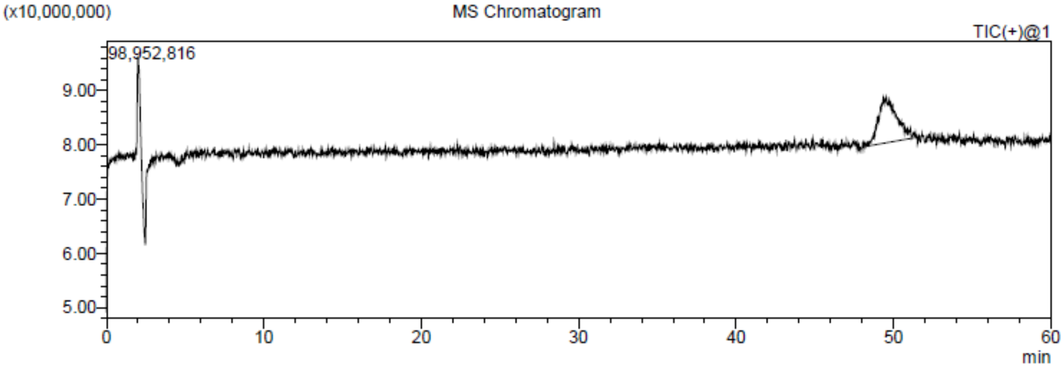


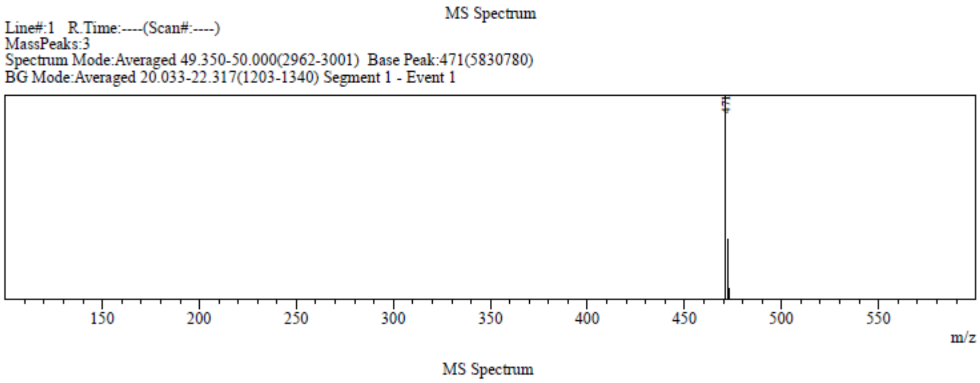


**Figure S10**. Determination of the enantiomeric excess of compound **4** (VUF13744, *S*-enantiomer), measured with SFC/MS (*System 3*) and detection at 254 nm.

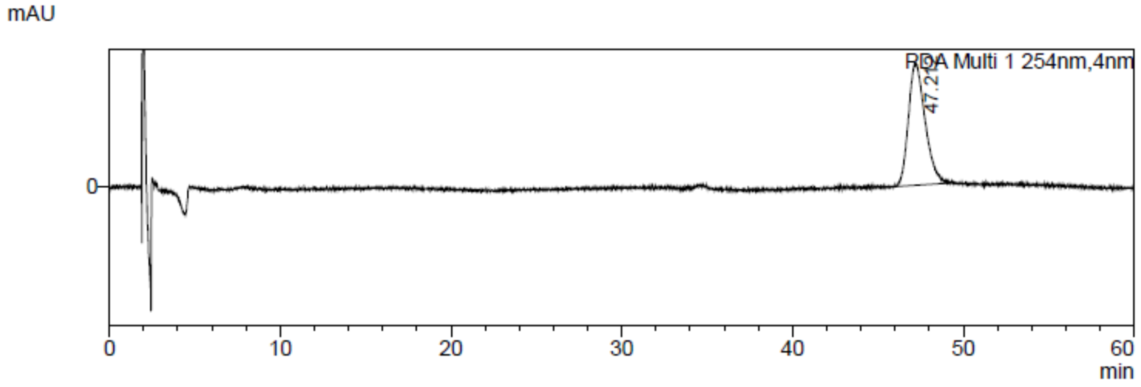


Peak Table

PDA Ch1 254nm

| Peak# | Ret. Time | Area% |
|---|---|---|
| 1 | 47.212 | 100.000 |
| Total | | 100.000 |

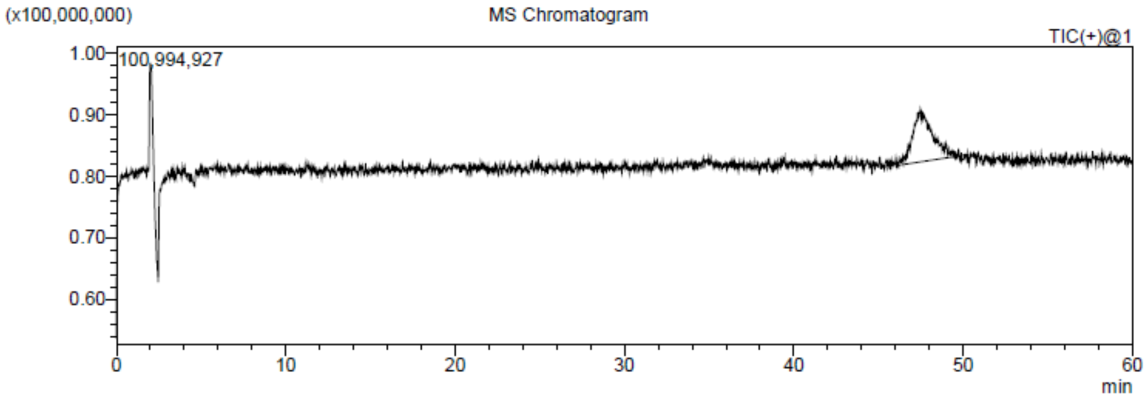


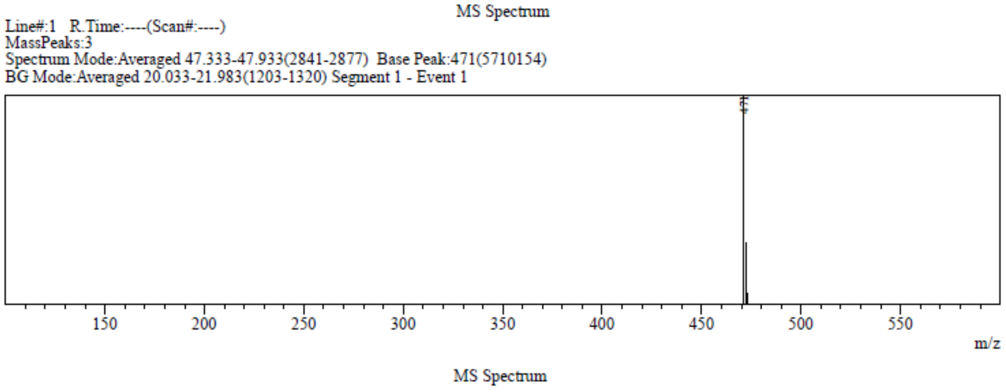


**Figure S11**. Determination of the enantiomeric excess of compound **5** (VUF15485, *R*-enantiomer), measured with SFC/MS (*System 3*) and detection at 254 nm.

**A**

Chromatogram: ³H

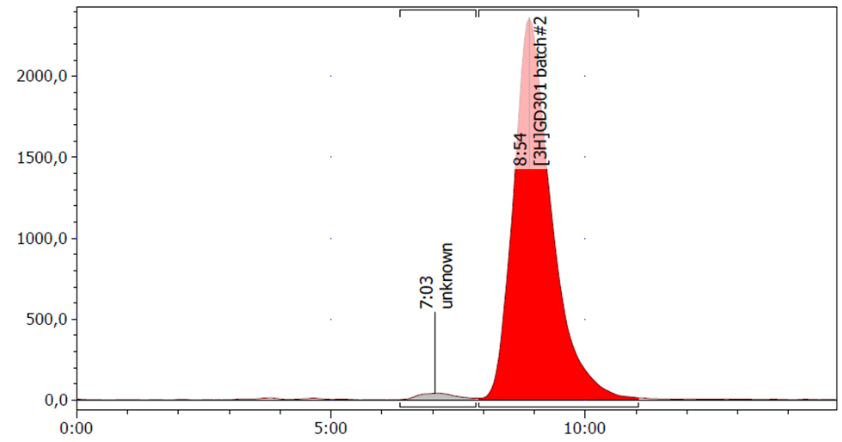


Regions: ³H Detector: ß-RAM

| Name | Start (mm:ss) | End (mm:ss) | Retention (mm:ss) | Height (Counts) | Area (Counts) | %ROI (%) |
|---|---|---|---|---|---|---|
| unknown | 6:21 | 7:51 | 7:03 | 51,0 | 767,0 | 1,84 |
| [3H]GD301 batch#2 | 7:54 | 11:03 | 8:54 | 2468,0 | 40987,0 | 98,16 |
| 2 Peaks | | | | | 41754,0 | 100,00 |

**B**

Chromatogram: UV

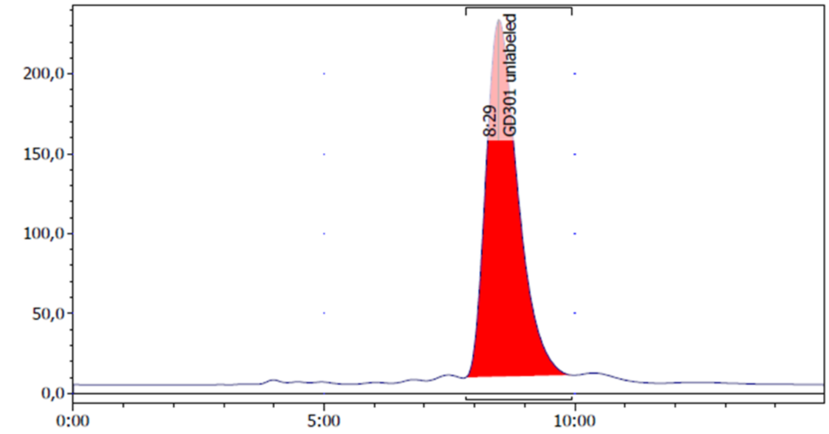


Regions: UV Detector:

| Name | Start (mm:ss) | End (mm:ss) | Retention (mm:ss) | Height (mAU) | Area (mAU·s) | %ROI (%) |
|---|---|---|---|---|---|---|
| GD301 unlabeled | 7:50 | 9:56 | 8:29 | 236,2 | 9913,6 | 100,00 |
| 1 Peak | | | | | 9913,6 | 100,00 |

**Figure S12**. Radiochemical purity of [$^3$H]VUF15485 (GD301). Chromatogram with (**A**) radiochemical detection and (**B**) UV detection of the unlabeled VUF15485 for peak identification. The time difference of 25 seconds for the peaks in panels **A** and **B** is in line with the detectors mounted in series.
